# The Brain/MINDS 3D Digital Marmoset Brain Atlas Version 2.0: Population-Based Cortical Parcellations With Multi-Modal Standard Templates

**DOI:** 10.1101/2025.08.28.672987

**Authors:** Rui Gong, Noritaka Ichinohe, Hiroshi Abe, Toshiki Tani, Mengkuan Lin, Takuto Okuno, Ken Nakae, Junichi Hata, Shin Ishii, Patrice Delmas, Shahrokh Heidari, Jiaxuan Wang, Tetsuo Yamamori, Hideyuki Okano, Alexander Woodward

## Abstract

We present our new Brain/MINDS 3D digital marmoset brain atlas version 2.0 (BMA2.0), a population-based 3D digital brain atlas of the common marmoset (Callithrix jacchus), designed to overcome the limitations of previous single subject atlases that are prone to structural biases arising from individual variation. Here, manually delineated cortical regions from 10 myelin-stained brains were used to create a generalized cortical parcellation. Newly refined subcortical regions from a previous atlas and a completely new cerebellum parcellation were also incorporated, resulting in a comprehensive whole brain parcellation for both hemispheres. To facilitate multimodal data analysis, the atlas package includes co-registered average templates for myelin and Nissl staining from the same individuals, ex vivo MRI T2 (91 individuals), and in vivo MRI T2 (446 individuals). Cortical flat maps and pial, cortical mid-thickness, and white matter surfaces are also provided. BMA2.0 provides a central brain space for multimodal data integration, spatial analysis, and comparative neuroscience. Standard formats and transformations are provided for easy integration into existing workflows and interoperability with existing atlases.

## Background & Summary

The Brain Mapping by Integrated Neurotechnologies for Disease Studies (Brain/MINDS) project^1^ was launched in 2014 by the Japan Agency for Medical Research and Development (AMED) in collaboration with an extensive network of research institutions. Using the common marmoset (Callithrix jacchus) as a model animal, the Brain/MINDS project was a decade-long endeavor that sought to deepen our understanding of the brain and develop new diagnostic tools and treatments for brain-related illnesses. Following the conclusion of the Brain/MINDS project in fiscal year 2023, a new project, Multidisciplinary Frontier Brain and Neuroscience Discoveries (Brain/MINDS 2.0) was launched in fiscal year 2024^2^. As data from a wide array of experiments is generated from such projects, a 3D brain atlas with spatial templates in a standard space is critical for data integration and analysis. This enables researchers to locate and identify brain regions and study the neural circuitry, connectivity, and function of these regions^3,4^.

We therefore present our new Brain/MINDS digital 3D marmoset brain atlas version 2.0 (BMA2.0), a fully symmetrical, population-based atlas with ex vivo and in vivo T2-Weighted Imaging (T2WI) MRI, myelin, and Nissl co-registered average brain templates. The multimodal templates simplify the process of data registration to the atlas, as mapping between different modalities can be challenging. This atlas aims to provide a more realistic representation of brain regions within the common marmoset population compared to existing atlases based on a single brain. Due to the amount of work needed to construct even a single brain atlas, a population-based atlas is considered the ‘Holy Grail’ of brain atlases. Creating a population-based marmoset brain atlas is a challenging task, as it involves integrating information from various imaging modalities and different histological techniques. We leveraged various AI-based computational methods in our atlas construction to make some of these tasks more manageable and improve image registration accuracy.

We used manually defined cortical delineations from 10 individual atlases, which collectively delineate 117 cortical regions per hemisphere. We employed a streamline-based methodology to assign the most probable label to each streamline within the cerebral cortex, while accounting for both the left and right hemispheres. This is combined with refined subcortical regions based on our previous BMA2017 atlas (156 regions) and a new cerebellum region atlas (45 regions), resulting in a total of 323 regions per hemisphere, providing the most comprehensive brain region coverage to date. This population-based atlas aims to capture the diversity of a population’s brains and give a more average shape of each cortical brain region by incorporating 10 individual brain atlases. This enables a more comprehensive understanding of the common marmoset’s brain, its organization, and its functions.

While previous atlases have focused on a single imaging modality or contrast, BMA2.0 uniquely integrates both Nissl and myelin staining within a common 3D coordinate space. These two histological modalities offer complementary laminar signatures: Nissl reveals cytoarchitectonic features based on neuronal cell body distribution. At the same time, myelin highlights fiber architecture through laminar patterns such as the middle band. This dual modality integration allows cross-validation of laminar features and supports population-level morphometric analyses. Furthermore, it establishes a robust anatomical framework for multimodal registration, facilitating comparison with in vivo MRI, tracer-based connectivity data, and spatially resolved molecular techniques such as spatial transcriptomics. Our population-based marmoset brain atlas is mapped onto a new symmetrical T2WI ex vivo MRI population average template based on 91 individuals from the eNA91 dataset. We have also created Myelin and Nissl contrast average templates based on histological data from those 10 individuals, as well as a new in vivo T2WI population average template derived from 446 individuals in the NA216 dataset, which is also mapped onto our new MRI average template space. This provides researchers with different modalities for registering, analyzing, and comparing their data using our new atlas. For interoperability, we also offered transforms from the BMA2.0 space to our previous releases, BMA2017 and BMA2019, as well as other common marmoset brain atlases, allowing analysis conducted within these releases to be easily transferred. Of note, we include the white matter parcellations of Liu et al.^5^ mapped to BMA2.0, thus providing full coverage of both gray and white matter structures in a single space.

The common marmoset is a small New World monkey that has come to be regarded as a valuable model animal due to its close phylogenetic relationship to humans^6,7^, its relatively small size, and its short gestation period. It is now used extensively in scientific research, particularly in studies related to neuroscience, including but not limited to genetics^8^, and brain-related diseases such as Alzheimer’s^9–11^, neuro degeneration^12,13^, and reproductive biology^14,15^. As neuroscience research continues to explore the complexities of brain structure and function, the development of a comprehensive brain atlas for the common marmoset becomes very important.

In prior work, the Brain/MINDS digital marmoset brain atlas 2017 (BMA2017)^16^ and the Brain/MINDS digital marmoset brain atlas 2019 (BMA2019)^17^ were developed. These atlases are available to the public through the Brain/MINDS Data Portal^1^. BMA2017 was based on a single brain^18,19^, where regions were annotated in the left hemisphere using a Nissl contrast and flipped across the mid-sagittal plane to give a complete annotation. The Nissl data was aligned, stacked, and co-registered with an ex vivo T2WI contrast of the same individual. For BMA2019, a population average T2WI ex vivo brain template was combined with the BMA2017 regions, along with the addition of cortical flat maps and cortical mid-surface data. Both atlases have been used in several influential marmoset brain studies, including the prefrontal cortex circuits study by Watakabe et al.^20^, and the Brain/MINDS NA216 (in vivo) and eNA91 (ex vivo) MRI database by Hata et al.^21^. While those two releases have improved our understanding of the structures of the marmoset brain, the inherent structural difference across individual brains, reflected in an atlas based on a single individual, may not provide the most accurate representation of different brain regions, as brains within a population can vary in size, shape, and anatomical structure.

Other well-known marmoset brain atlases exist, such as the population-based cortical template by Majka et al.^22^, that provides a similar population-based cortical atlas, but this uses Nissl-based cortical region delineations for the left hemisphere only, whereas we provide cortical and subcortical delineations for both hemispheres, and provide Nissl and myelin templates from the same individuals of the whole brain. We also include ex vivo and in vivo T2WI MRI templates, making it suitable for MRI-based studies. The marmoset brain mapping atlas (MBM), version 1 by Liu et al.^23^, version 2 by Liu et al.^5^, version 3 by Liu et al^24^, version 4 by Tian et al.^25^, and version 5 by Zhu et al.^26^, Hao et al.^27^ provides a comprehensive collection atlases of the marmoset brain. Each new version improved anatomical precision, multimodal integration, and utility for cross-subject and cross-species neuroimaging studies. The marmoset brain mapping atlas version 3 (MBMv3) by Liu et al.^24^ offers a valuable template by integrating two existing brain parcellations, the marmoset brain mapping atlas version 1^23^ and our BMA2017^16^. However, its application is constrained by the use of only two brain parcellations, a limitation that hinders the sufficient diversification necessary to mitigate potential biases associated with individual variability. Saleem et al.^28^ presented the Subcortical Atlas of the Marmoset (SAM), a high-resolution 3D digital atlas that delineates 251 subcortical structures for the whole brain. It includes 211 gray matter regions and 40 white matter tracts. The atlas was constructed using a single brain and integrating ultra-high-resolution ex vivo MRI with histological validation, enabling the precise segmentation of deep brain regions.

Beyond serving as a reference framework, the multimodal nature of our dataset also enables morphometric studies. In particular, as part of the technical validation of our atlas, we leveraged the laminar contrast observed in myelin-stained sections to quantify the variation in thickness of layers 1-3 across the cerebral cortex. While high magnification Nissl staining is often used to define cortical layers, it may be insufficient at lower resolutions due to the variable cell density in layer 4 and the intense staining in large layer 5 neurons. In contrast, the myelin-stained middle band provides a more consistent and reliable laminar signature at the resolution used for population-based mapping. Using this signal, we observed systematic dorsal-ventral and rostral-caudal gradients in supragranular thickness, highlighting the biological richness of our atlas and its potential for comparative cortical studies.

Our new multimodal, population-based marmoset brain atlas serves as a state-of-the-art reference for brain researchers. It represents a significant advancement for our understanding of marmoset brain structure and function. BMA2.0 is expected to advance the field of neuroscience while simultaneously offering insights with potential implications for human brain health and disease research.

## Methods

### Data acquisition and histological annotation

#### Brain slicing and staining protocols

Each ex vivo brain was initially placed in dry ice, after which coronal brain sections were gathered, rostral to caudal, at a thickness of 50 µm. Sections were assigned in a repeating sequence for different histological processing in the following order: myelin staining, Nissl staining, and tracer imaging, where the tracer images were used in separate studies. This alternating scheme (myelin-Nissl-tracer) ensured that the histological series were evenly spaced at regular intervals throughout the brain, enabling multimodal analysis across comparable anatomical levels.

Concurrently, during the sectioning process, block face images were taken after each coronal section using a camera mounted at a fixed position from above. The total number of block face images produced by each brain was approximately 600. More details are provided in Table 1.

**Table 1.**
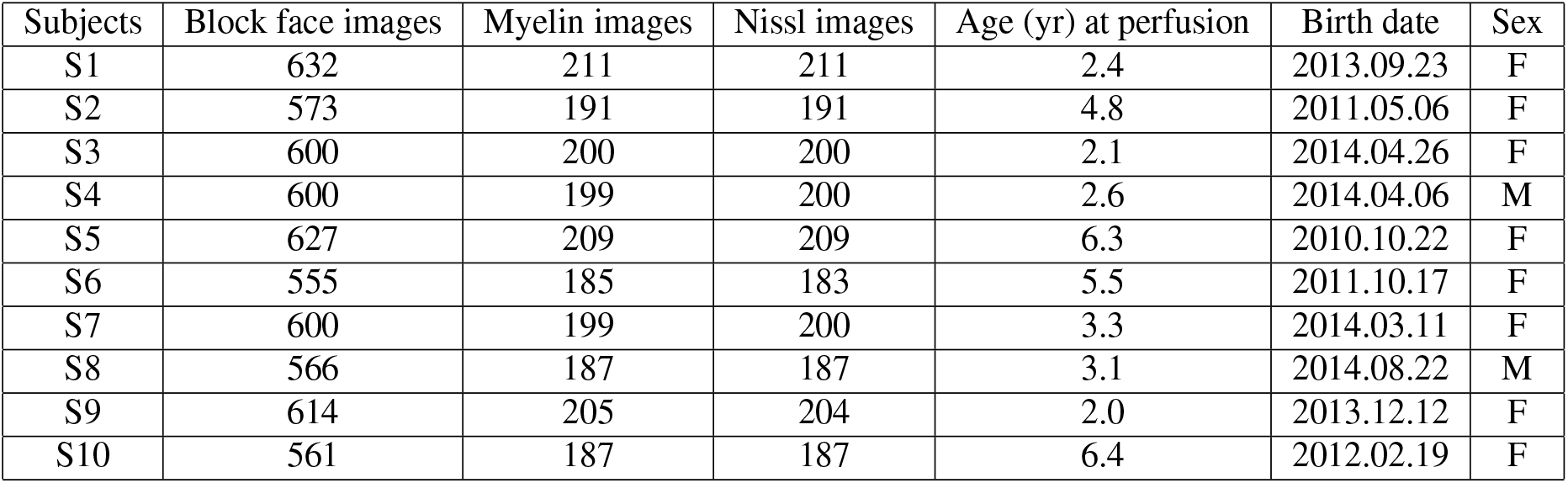
Information on the images to be processed and registered for population-based cortical parcellation, average myelin-stained and average Nissl-stained template construction.

A NanoZoomer 2.0-HT batch slide scanner (Hamamatsu Photonics) was used to scan each section at high resolution ( ≈0.46 µm/pixel). A more detailed explanation is provided in Abe et al.^29^. Data on the 10 marmosets used in this work are listed in Table 1.

#### Ex vivo and in vivo MRI acquisition

Ex vivo T2WI MRI scans of the 10 adult marmosets were acquired using a 9.4-T BioSpec 94/30 unit (Bruker Optik GmbH, Ettlingen, Germany) and a transmit and receive solenoid type coil with an inner diameter of 28 mm.

In vivo T2WI MRI scans were obtained as part of the NA216 study. These data contains in vivo multimodal MRI data from 216 common marmosets aging between 0.8–10.3 years. Animals were anesthetized and scanned with the same 9.4-T BioSpec 94/30 unit, and a transmit and receive coil with an inner diameter of 86 mm. A more detailed description of the data acquisition process can be found in Hata et al.^21^.

#### Manual delineation of cortical regions on myelin-stained slices

Cortical region delineations were identified by neuroanatomists and drawn onto every individual myelin-stained tissue slice (150 microns between each myelin-stained slice) in both hemispheres, for 10 individuals. Two histological techniques were used to help identify brain regions, myelin staining (fiber tracts) and Nissl staining (cell bodies), in conjunction with the reference border definitions and nomenclature provided by Paxinos’ marmoset brain atlas^3^.

#### Manual background segmentation

The background segmentation, which is used to separate brain tissue from background, was manually drawn for all myelin (1/3 of block face slices) and block face slices (roughly 600 slices) of the 10 brains. This segmentation information was later used in our processing pipeline in two ways: first, as additional constraints for image registration, and second, as training data for artificial intelligence (AI) algorithms for image segmentation tasks.

#### Convert cortical region delineations to machine-readable format

Myelin images, along with their corresponding cortical region delineations, were first processed using a module of our NanoZoomer image processing pipeline (NZP)^17^, which converts region delineation drawings (in SVG format) into 2D TIFF images with their respective region labels. Example outputs from our pipeline are shown in Figure 2B and Figure 2C. 16-bit grayscale values were assigned to each region, representing its region number using a region lookup table as defined in the BMA2017 and BMA2019 releases.

**Figure 1.**
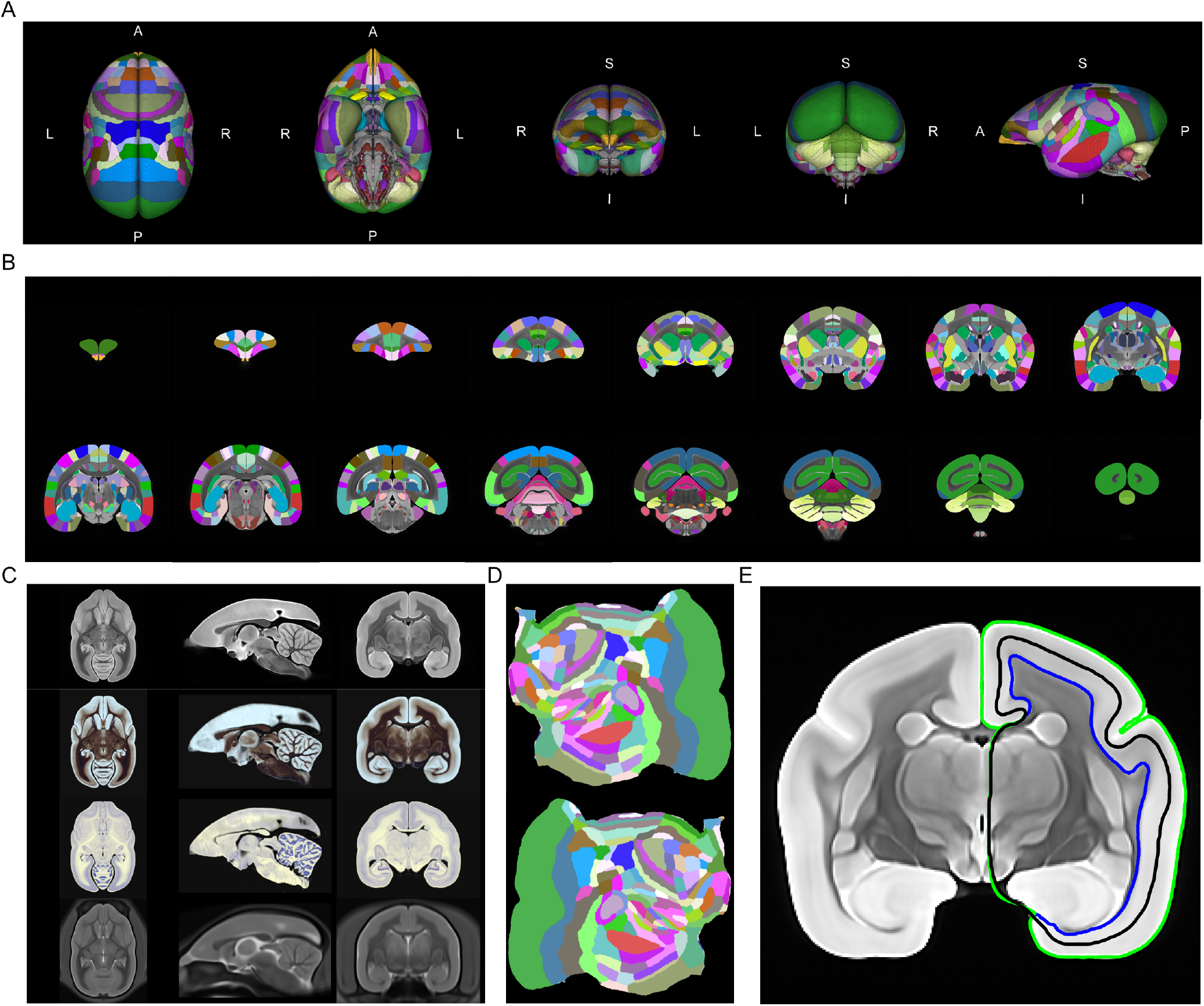
Overview the BMA2.0 package. (A) 3D visualization of the BMA2.0 atlas, from left to right: superior, inferior, anterior, posterior, and left hemisphere views, respectively. (B) Coronal views of the BMA2.0 population-based atlas (117 cortical regions, 156 subcortical regions, and 45 cerebellum regions per hemisphere) from anterior to posterior with 40 slice intervals, on top of the BMA2.0 population average T2WI MRI template. (C) Axial, sagittal, and coronal views of the BMA2.0 population average T2WI MRI (*N* = 91), myelin (*N* = 10), Nissl (*N* = 10), and in vivo T2WI (*N* = 455) templates. (D) From top to bottom: left and right hemisphere cortical flat maps with BMA2.0 atlas label mapping. (E) Example 2D coronal view showing left hemisphere cortical mid-thickness (black), pial (lime green), and white matter (blue) surfaces (as lines) mapped onto the BMA2.0 population average T2WI MRI template.

**Figure 2.**
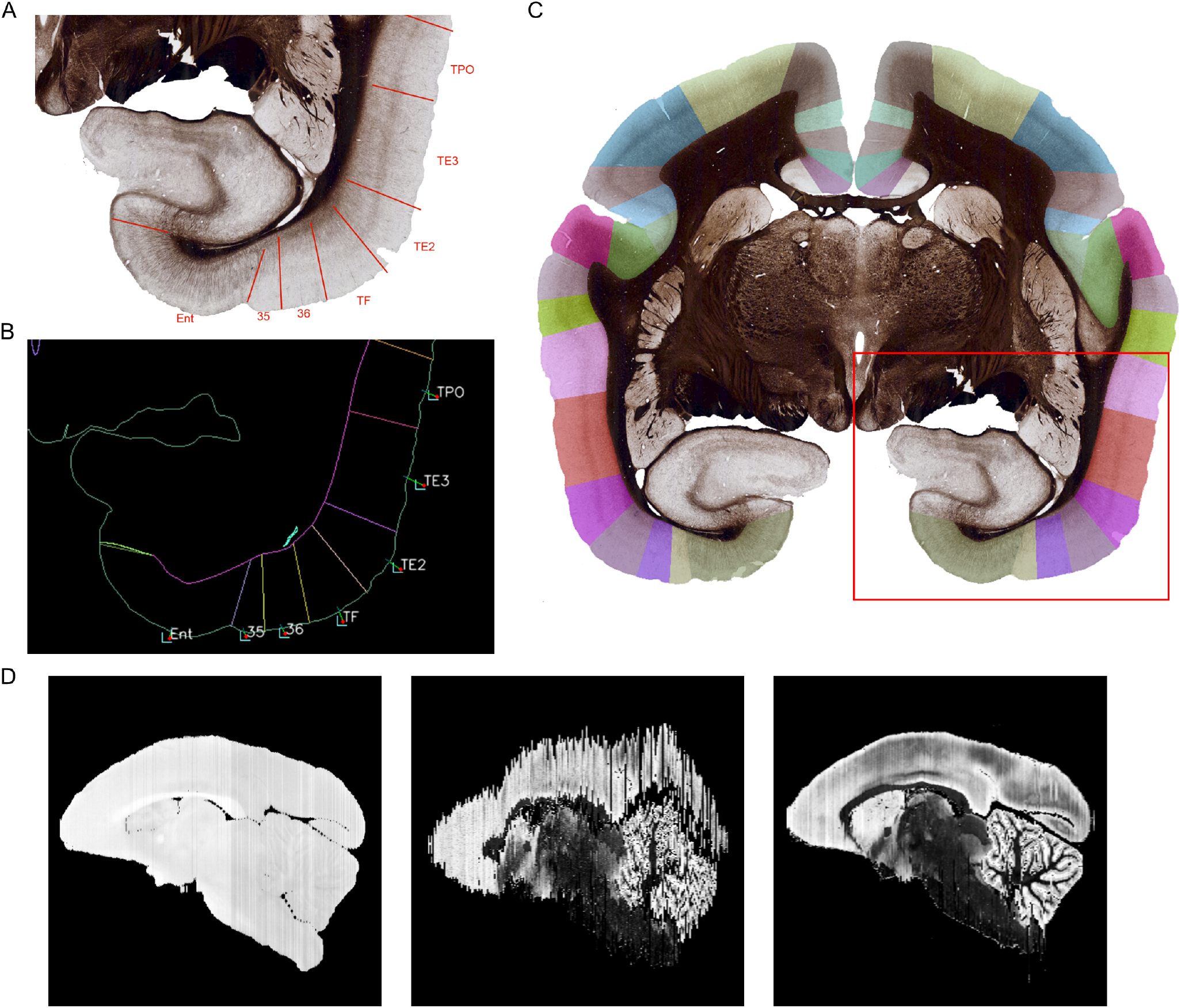
Example of NanoZoomer pipeline image conversion and image registration. (A) Inset of the red subregion shown in C, showing cortical region delineations identified by neuroanatomists in myelin histology. (B) Converted machine-readable region delineations of A using the Nanozoomer image process pipeline. (C) Converted region delineations overlay onto the original myelin image. (D) Example of image registration of myelin to block face in sagittal view, comparing myelin volume (image stacking) before and after image registration. From left to right: block face volume, myelin volume before image registration, and myelin volume aligned to block face volume after image registration.

Following the machine conversion of the region delineations for all the slices, manual reviews were conducted to identify and rectify any errors that occurred during the conversion process. We identified conversion errors, including incorrect region assignments, multiple region assignments to the same regions, and unenclosed region borders. Such problems were fixed manually and then reprocessed using our pipeline. In instances where region delineations were absent due to damaged or missing tissue, the missing slice was replaced by a duplicate of the previous slice. This manual checking process ensured the correct region delineation conversions for all slices (approximately 200 per brain) across all 10 brains.

### Image registration pipeline

We developed a workflow to carry out 2D and 3D image registration accurately, leveraging AI-based image processing techniques to enhance registration accuracy. This allowed us to transform 2D sections into complete 3D brains, use non-linear mapping to create an average brain shape, and map all data into this standardized space.

#### 2D and 3D registration using Advanced Normalization Tools (ANTs)

Image registration is a challenging problem in medical imaging. The objective involves aligning images, whether of the same or different modalities, into a reference coordinate system. This is achieved through linear and nonlinear image warping techniques, which establish a spatial correspondence between the pixels or features present in the images. This is one of the most essential processes of digital brain atlasing, as we need to restore histological data, including myelin and Nissl, to their original (block face) shape using 2D image registration. 3D image registration was also used to transform between an individual block face volume and ex vivo MRI volume, as well as to map all 10 individual brain atlases into a common standard brain template. As shown in Figure 2D, stacking myelin-stained images without image registration results in an unaligned brain volume, compared to its original block face volume, and this is restored through precise image registration.

For our work, we used Advanced Normalization Tools (ANTs) version 2.4.4 (“Eotapinoma”, released May 2nd, 2022) to perform both 2D image and 3D volume image registrations (with linear and non-linear deformations) ^30,31^. We mostly used two registration shell scripts, *antsRegistrationQuick*.*sh* and *antsRegistrationSyN*.*sh*^2^, which were executed under the Ubuntu version 22.04 operating system.

#### AI-based image processing techniques for improving image registration

We used Artificial Intelligence (AI) algorithms to enhance the accuracy of image registration across different image modalities (myelin, Nissl, T2WI MRI, and block face) by either providing additional information, such as cortical boundary constraints, or translating images into similar modalities. In image registration, additional segmentation constraints function as landmarks on the images, providing the registration algorithm with information on the areas and regions that should be matched.

#### Image translation of Nissl to synthetic myelin

Due to the distinctiveness of color and contrast between myelin and Nissl images, it is very challenging to perform image registration that accurately aligns internal brain structures. To improve the accuracy, we trained a CycleGAN model, a type of generative adversarial network popular for unpaired image-to-image translation, to translate Nissl images into myelin-like color and contrast ^32^. The main benefit of this algorithm is that it eliminates the need for image pairing during the training process, thereby simplifying the preparation of training images for the algorithm.

To prepare the training data for CycleGAN, we used myelin and Nissl images from our 10 marmoset brains (around 200 myelin and Nissl images per individual). The original myelin images were resized to align with the dimensions of the background segmentation images, as the original myelin images were acquired at extremely high resolution using a NanoZoomer 2.0-HT scanner (20× objective, 455 nm/pixel as detailed in Tani et al.^33^), and the manual segmentation was drawn under significantly lower resolution.

Background segmentation was then applied to myelin images to remove background noise, such as liquid and parts of brain tissue from other slices. Manual background segmentation was not performed on the Nissl images as brain region delinations are drawn onto myelin images, so the training dataset for Nissl included background artifacts, such as fragments of brain tissue from adjacent slices, liquid residual, etc., in some images.

Next, we applied padding to each image in the training set to ensure that all images were the same square size. For each brain, we used either the maximum width or height (*σ*_*max*_) from all the myelin images, to keep the relative physical size for all images. Zero padding was applied to pad each image into a square size, with a new length and height = *σ*_*max*_, while keeping the brain tissue at the center. Finally, all padded images were resized to 512 × 512 pixels. The same padding process was used for the Nissl images.

Image augmentation was employed to increase the number of training images in our limited dataset thereby increasing robustness and avoiding overfitting^34^. We used the Python package Argumentor^35^ to do this.

Zooming and rotation augmentations were applied to the myelin and Nissl training images. In total, we used 9261 myelin and Nissl images to train the network. Figure 4A shows some examples of Nissl images translated to synthetic myelin images using our trained CycleGAN network.

**Figure 3.**
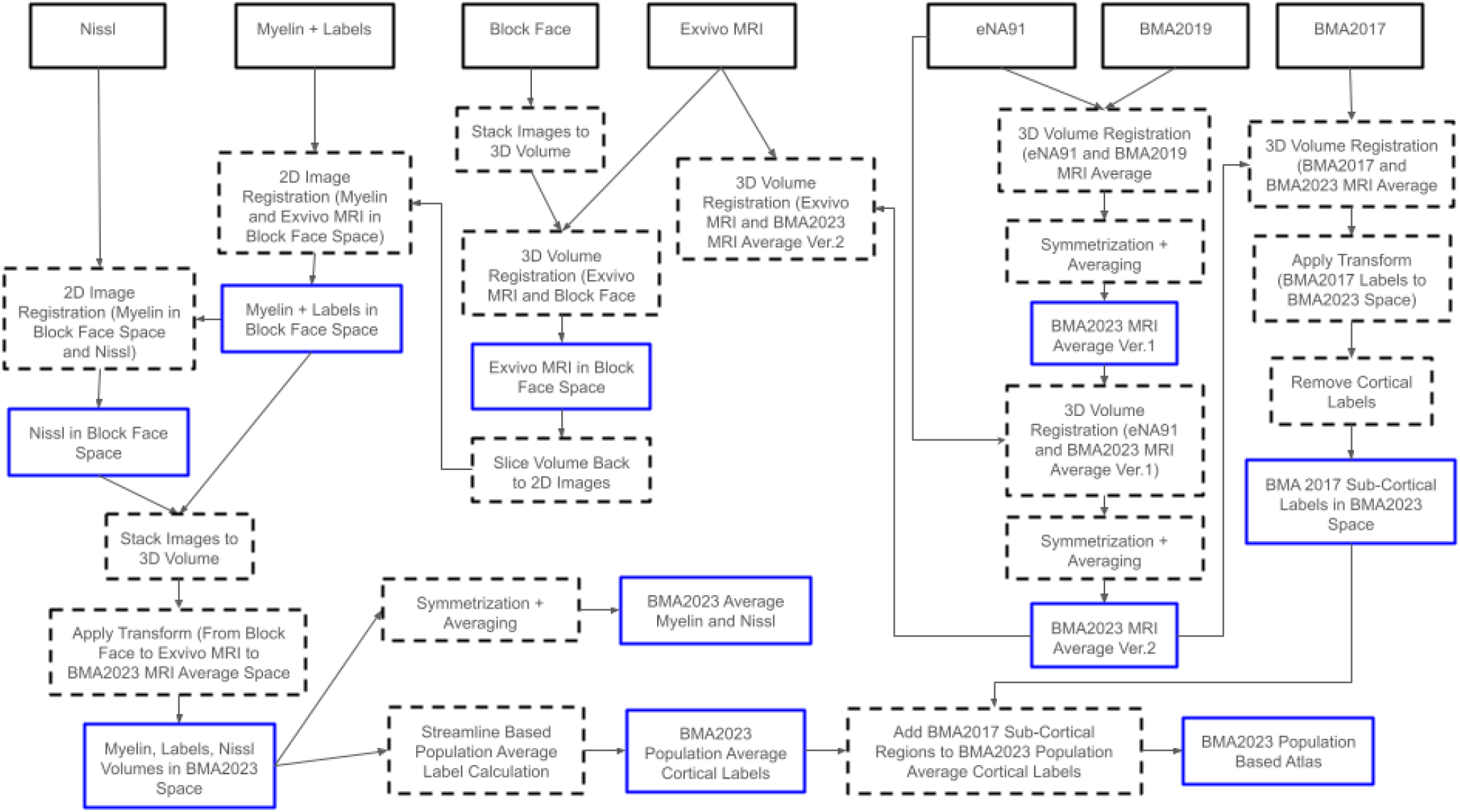
Overview of the BMA2.0 population-based atlas and average templates construction. Black boxes indicate the original data sources, dotted boxes indicate image processing applied, and blue boxes indicate outputs.

**Figure 4.**
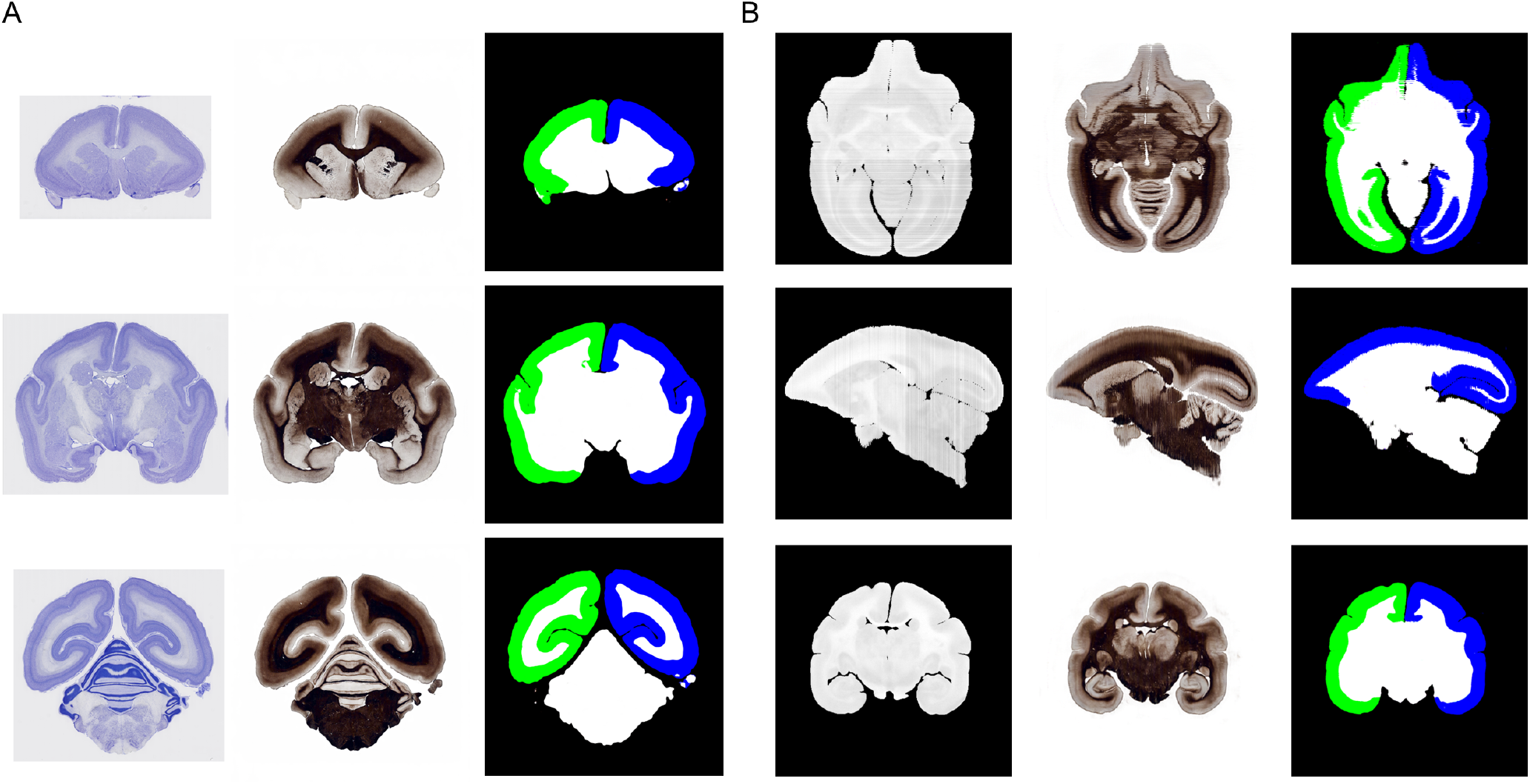
Example outputs of image translation using CycleGAN and cortical segmentation using Pix2Pix (A) From left to right: examples of synthetic myelin images generated from Nissl using CycleGAN. Then, from synthetic myelin images, cortical segmentation was generated using Pix2Pix’. (B) From left to right: an example of a whole synthetic myelin volume generated from images from a block face volume using CycleGAN, followed by a cortical segmentation volume generated from the synthetic myelin volume using Pix2Pix. Blue, green, white, and black represent the left cerebral cortex, the right cerebral cortex, non-cortical areas, and the background, respectively, in the axial, sagittal, and coronal views (from top to bottom) for figures A and B.

#### Cortical boundary segmentation of translated Nissl

To further improve the accuracy of image registration, we can include additional constraints, such as segmentation, in conjunction with the image data^36^. Segmentation provides additional information to help identify general regions to be matched during image registration. In the case of Nissl to myelin image registration, we would like to include segmentation masks that identify several brain regions: cortical regions (left and right), subcortical regions, and background, to guide the matching of relevant brain regions during the registration process..

To create the cortical boundary segmentation images, we employed Pix2Pix, a GAN-based algorithm first introduced by Isola et al.^37^, to train and generate the left/right cortical boundary and subcortical segmentation for myelin contrast images. Compared to CycleGAN, Pix2Pix learns a mapping from an input image to an output image based on paired examples, meaning Pix2Pix requires the preparation of training images by pairing each myelin image with its corresponding segmented mask.

To prepare training segmented mask data, we first generated cortical and non-cortical brain regions segmentation from cortical region delineation data generated by our NanoZoomer pipeline, and then combined them with background segmentation data. To generate left/right hemisphere cortical boundary segmentations from cortical region delineations, we first identified the center of an image and then assigned either the left or right hemisphere to each cortical segmentation based on its location. For all slices, we manually corrected any incorrect assignments in the output. Next, we combined the left/right cortical segmentations with the background segmentation information to identify the non-cortical brain region. We assigned four color codes: blue, green, white, and black, representing the left hemisphere cortical regions, the right hemisphere cortical regions, non-cortical areas, and the background, respectively.

We paired the original myelin images with the cortical segmentation images (from 6 brains, totaling 1181 image pairs). We used Argumentor^35^ to apply rotations and zooming to both myelin and segmentation image pairs. A total of 8192 image pairs were used to train the Pix2Pix network.

We used the trained network to generate cortical boundary segmentation using synthetic myelin images derived from Nissl images, as shown in Figure 4A, providing additional segmentation constraints to improve image registration from Nissl to myelin.

#### Background removal in Nissl images

To further improve the image registration accuracy of Nissl to myelin images, we separated brain tissue from the background, as the background contains noise such as liquid or brain tissue from adjacent slices, which may affect registration accuracy at the brain section boundary. However, unlike myelin, manual background segmentation was not available for the Nissl images.

To generate the background segmentation for Nissl, we took advantage of the myelin image training set for CycleGAN, which already incorporates background segmentation into the images. This means that our trained CycleGAN network, using this training set, can generate synthetic myelin images with clean backgrounds from Nissl image input. Figure 4 shows examples of synthetic myelin images generated from Nissl images.

From these new images, the background was easily segmented using a threshold technique, where pixels corresponding to white or near white intensities were treated as background. While this approach effectively addressed the removal of the background, it did not automatically eliminate extra tissue sometimes present in the Nissl images. For such images, we manually removed any extra brain tissue.

#### Image translation of block face to synthetic myelin

To enhance the accuracy of image registration from myelin to block face images, we again used CycleGAN, similar to the Nissl to synthetic myelin image translation. To prepare training images, we first applied background segmentation to remove background noise in the myelin images and the block face images. Image padding was then applied to all images for each of the 10 brains. These images are then used to train the CycleGAN network.

The lack of texture in the block face images meant that some structures, such as in the cerebellum and the depth of cortical layer patterns, were unrealistic in the generated synthetic myelin. We have also noticed that synthetic myelin images from block face picking up features such as tissue shrinkage, thus modifying the overall tissue shape. Similar issues were not observed in Nissl to synthetic myelin translation, likely due to tissue shrinkage that was present in both Nissl and myelin images. Therefore, we did not use the synthetic myelin directly for image registration, as such results may warp the cerebellum, the cerebral cortex layers, and overall brain shape away from the original block face data in unexpected ways. However, the synthetic myelin from block face was used to generate cortical and non-cortical segmentations, as described in the next section, which were then used to guide 3D volume registration of individual MRI contrasts to block face volumes.

#### Cortical boundary segmentation of translated block face

We used the trained Pix2Pix network to generate cortical segmentations for each slice of synthetic myelin from block face data using CycleGAN. Segmentations were then converted to a volumetric format for each of the 10 brains in block face space. Figure 4B shows an example of the result. The volumetric segmentation separates cortical regions, non-cortical regions, and the background for each brain. These were used as markers to guide the non-linear 3D registration of individual MRI volumes to individual block face volumes. This ensures the alignment of brain structure segmentation between the block face and MRI volumes for each brain.

### Brain region parcellations and templates construction

#### Population average ex vivo T2WI MRI brain template construction

We created a new population average ex vivo MRI T2WI brain template using the Brain/MINDS eNA91 dataset (Hata et al.^21^). eNA91 contains ex vivo MRI scans of 91 individuals with an average age of 5.27 *±* 2.39 years. We used the T2WI MRI scans for the template construction. Our template is symmetric across the left and right hemispheres, allowing it to serve as an unbiased basis for analysis. This high-quality average template was used as the standard space for our atlas.

All 91 scans were first registered to the BMA2019 population average T2WI MRI template (volume size 300 × 360 × 220 voxels at a resolution of 100 µm) using ANTs. This gave us 91 individual brains in the BMA2019 space. We next made each registered brain in the eNA91 dataset left/right symmetric^3^. To achieve this, we flipped each brain volume across the midsagittal plane and then registered it with the original one. Halfway transformations in both forward and backward directions were then calculated and applied to the flipped and original brain volumes, transforming them into almost symmetric versions. Lastly, the average of the symmetrized, flipped, and original brains was used to create each left/right symmetric brain.

The first iteration of the average template was created by averaging the 91 symmetric brains, with a final flip of the left hemisphere across the midsagittal axis to obtain a voxel-level symmetric population average template.

To further refine the BMA2.0 population average template, we re-registered the 91 symmetric brains to the first iteration of the template and repeated the averaging process. This gave us a slightly more refined version of the BMA2.0 population average template. Areas such as the lateral sulcus were improved compared to the first iteration. This refined version was used as the new standard space for our BMA2.0 dataset. We used the same (Talairach-like) RAS anatomical coordinate system as in our previous atlases (BMA2017 and BMA2019). RAS is defined as the X-axis increasing in value from Left to (R)ight, Y from Posterior to (A)nterior and Z from Inferior to (S)uperior. The resolution of our new BMA2.0 ex vivo T2WI MRI template is the same as BMA2019, 300 *×* 360 *×* 220 pixels, with a voxel size of 100*µ*m.

#### Individual and Average MRI cortical boundary segmentation and registration

To accurately transform data from the 10 brains with region labels, myelin, and Nissl to the new BMA2.0 space, it is preferable to perform registration between data with the same or similar modality. Since the standard space of BMA2.0 was constructed from ex vivo T2-weighted imaging (T2WI) MRI data, we first registered the 10 individual T2WI MRI volumes to this space. For accurate registration, we created cortical boundary segmentations as constraints to ensure precise alignment..

Manual segmentation was previously performed on BMA2019 to segment the left and right hemispheres of the cerebral cortex, white matter, subcortical structures, and cerebellum. We used ANTs to transform these segmentations to the 10 individuals. We manually checked and refined the transformed region segmentations for each of the 10 individual brains using the segmentation editor module from 3D Slicer^38^ (version 5.2.2 was used)^4^ to ensure they reflected the nuances of each brain. The same process was repeated to create a cortical boundary segmentation for BMA2.0.

We then used ANTs to non-linearly register each of the 10 individual MRI volumes to the BMA2.0 average template, with the cortical segmentations providing additional alignment constraints. The resulting transforms allow us to accurately transform data between the individual MRI space and the BMA2.0 average template space.

#### Iterative image registration of individual myelin to individual block face

The transformation from individual myelin/region labels to the original brain shape is needed to go from 2D slices to a 3D volume. We first obtained accurate registrations between individual MRI volumes and individual block face volumes by stacking individual block face images and their corresponding cortical boundary segmentations into 3D volumes. Then, we registered them to individual MRI volumes with cortical boundary constraints.

Individual MRI volumes were then transformed to block face space, allowing us to create 2D MRI contrast images for each block face coronal slice, with the slice numbering matching the corresponding myelin and Nissl image data. Image registration of individual myelin images to corresponding 2D MRI T2WI images in block face space was then done. This approach was taken because myelin and MRI images exhibit similar textures in brain structures when compared to the relatively textureless block face images. Registered myelin images and their cortical segmentation were stacked into a 3D volume, giving a myelin volume (first iteration) and a segmentation volume in individual block face space.

To iteratively improve the reconstruction, we applied a 3D median image filter (with a voxel radius of 1) to the transformed myelin volume. A 3D median filter was chosen due to its ability to reduce noise while preserving edges in the image^39^. This step blurs the volume, reducing noise between slices in 3D (specifically in the cerebellum region) while preserving structural boundaries. The blurred myelin volume was sliced into coronal plane images, and registration of the original myelin images to the blurred myelin with a cortical segmentation constraint was done. As the images now have similar contrast, this step provided a much better registration of the internal structures in the myelin images while preserving 3D structural consistency. Again, we stack the registered myelin images into a 3D volume (registered myelin volume iteration 2).

This 3D myelin volume (iteration 2) was then registered to the MRI volume in block face space to generate a new transform. This transform was modified by zeroing out one dimension (AP-axis), so that slices are not shifted along this axis, and applied to the 3D myelin volume (iteration 2). This refines the alignment of internal structures and ensures that myelin slices align correctly in 3D. This process was carried out for another (third) iteration to give a final mapped 3D myelin volume in block face space.

#### Myelin average template construction

Previously calculated transforms from individual block face volumes to individual MRIs and individual MRIs to BMA2.0 were used to map individual myelin volumes into the BMA2.0 template space. To increase the total number of brains and make a symmetric template, we left/right flipped each of the 10 myelin brains to get a total number of 20 myelin brains. The myelin average template was created by averaging the data from the 20 myelin brains. The resolution of the BMA2.0 average myelin-stained brain is 300 *×* 360 *×* 220 pixels, with a voxel size of 100*µm×* 100*µm×* 100*µm*.

#### Nissl average template construction

To make the average Nissl-stained brain, we first registered synthetic myelin from Nissl to the myelin images in block face space, along with cortical segmentation constraints. The results were stacked to create a 3D Nissl volume in the block face space. The previously calculated transforms, which map individual block face space to individual MRIs, and then to the BMA2.0 population average space, were used to map data into the BMA2.0 space. We then flipped the 10 transformed Nissl brains to get a total of 20 brains and averaged them. This provided us with the first iteration of the average Nissl-stained brain.

To refine the result, we transformed the average Nissl-stained brain back to each block face space. This data was sliced into 2D coronal images, which were then registered with their corresponding original Nissl images to get a more precise alignment than before. Registered images were then stacked into 3D volumes and transformed to the BMA2.0 average space using the same transforms. Again, we flipped the 10 brains to obtain 20 brains and performed averaging to create a second iteration of the Nissl average template. Due to better alignment, the second iteration of the Nissl average appears less blurry with a more precise definition of layer 4 within the cerebral cortex, as well as a more distinct cerebellum. The resolution of the BMA2.0 average Nissl-stained brain is 300 *×* 360 *×* 220 pixels, with a voxel size of 100*µm×* 100*µm×* 100*µm*.

#### Population average in vivo T2WI MRI brain template construction

An in vivo MRI brain template was created to support resting-state fMRI and structural connectivity studies^40–42^. In addition, by providing an in vivo template, the process of skull stripping can be made easier or even unnecessary if the goal is to map the atlas to the data.

We created a new in vivo standard template based on our BMA2.0 space using data from the NA216 dataset^21^. The inclusion of the skull in in vivo data is the main challenge in registering it to our BMA2.0 average template. To overcome this, we took the existing in vivo NA216 average template (*N* = 216), made it left/right symmetric, and manually segmented the brain tissue to strip the skull using 3D Slicer’s segmentation tools. We then registered the skull stripped in vivo template to our BMA2.0 average template. The resulting transforms were then used to transform the in vivo average template with the skull into our BMA2.0 space.

We then took all the individual in vivo T2WI data (*N* = 455) from the dataset and registered it to the in vivo average template in the BMA2.0 space. Next, we took these transformed in vivo brains, flipped them across the midsagittal axis, and calculated the average to create our final BMA2.0 in vivo T2WI MRI average template.

#### Individual marmoset brain atlas construction

To construct population-based cortical parcellations, we first need to build the 10 individual cortical parcellations. Region label images for each 2D myelin image space are raster images where each pixel value corresponds to a region number listed in a brain region lookup table. The transforms from individual myelin to individual block face space were applied to the label map images. These were stacked to create individual 3D label map volumes. Coronal, sagittal, and axial views of the 10 individuals are shown in Figure 5. The data was then transformed to its individual MRI space, and subsequently to the BMA2.0 average template space. Nearest neighbor interpolation was used when transforming the labels.

**Figure 5.**
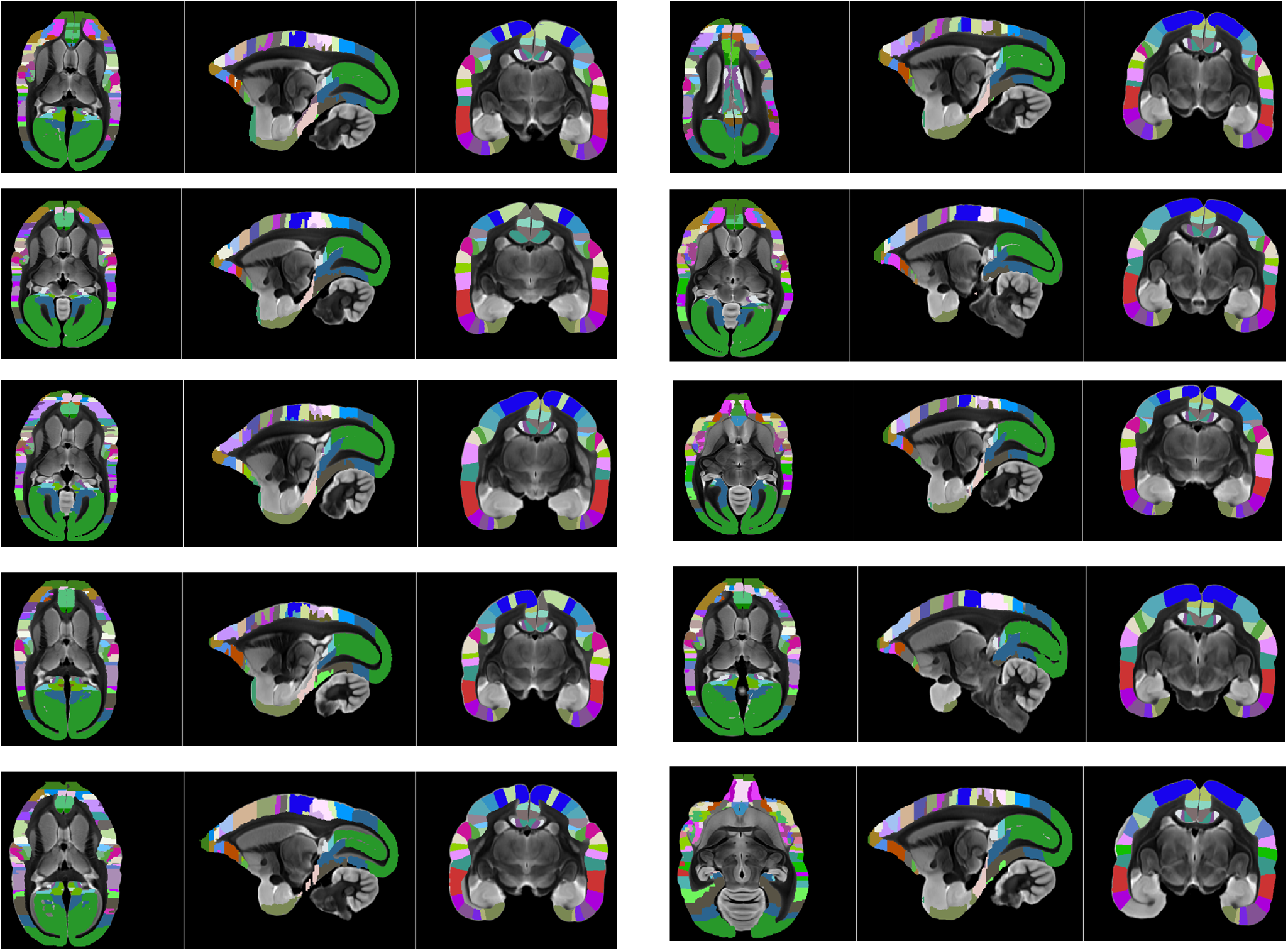
Axial, sagittal, and coronal views of the 10 individual cortical parcellations, in their individual shapes, on top of their individual ex vivo T2WI MRI contrasts. These brains were used for our BMA2.0 construction, where individual brain cortical parcellations were reconstructed using our NanoZoomer image processing pipeline. Variabilities between individual cortical parcellations are noticeable, highlighting the disadvantage of atlases based on a single brain. Brain annotations were made using coronal sections, highlighting the approach’s disadvantage, where cortical region boundaries may not align with the correct cortical columnar anisotropy (i.e., the direction normal to cortical layers). This is corrected using a streamline approach to cortical parcellation.

This process was applied to all 10 individual brain datasets, resulting in 10 distinct brain cortical parcellations in the BMA2.0 space. All 10 transformed cortical parcellations have a resolution of 300 *×* 360 *×* 220 pixels, and with a voxel size of 100*µm×* 100*µm×* 100*µm*.

We wanted to create region labels that are truly symmetric to accompany our new population average template. Similar to other modalities, we flipped the 10 individual cortical parcellations across the midsagittal axis to obtain a total of 20 cortical parcellations. This information was used to construct the final labeling at each voxel. Since the final atlas is left/right symmetric, we could reduce the amount of computation needed for cortical parcellation construction in one hemisphere only and then mirror the result.

#### Population-based cortical structure delineation

To calculate the population-based label in the BMA2.0 space using the 10 individual cortical parcellations, we used a streamline-based method to assign the most probable label per streamline flowing along the gradient of the solution to Laplace’s equation with two (Dirichlet) boundary conditions defined at the white matter and pial borders^22,43^. A streamline-based approach, using the optimally smooth solution of Laplace’s equation, is assumed to respect the cortical columnar direction and is employed to refine coronal section-based region parcellations mapped into 3D. Laplace’s equation is defined as:

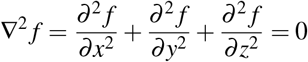

In our case, the solution *f* yields smoothly changing values as it transitions from the white matter to the pial boundary in 3D. To compute this solution, we first needed to distinguish the cortical mask, background, and inner cortical (white matter boundary) surface segmentation and define Dirichlet boundary conditions.

For the cortical mask and background segmentation, we used transforms from BMA2019 to BMA2.0 to convert the existing cortical mask and background segmentation for BMA2019 to BMA2.0. We then used 3D Slicer to make adjustments to the cortical mask in challenging areas such as the gap in the lateral sulcus. We then manually defined the inner cortical surface within the cortical mask for BMA2.0. The cortical mask and background segmentation information were used to delineate the pial surface of the cerebral cortex. This was achieved by examining the overlap between the outer surface of the cortical mask and the background, excluding the inner surface.

Once we have defined the pial and white matter surfaces, we can use them to solve Laplace’s equation within the cortical mask region. Here, we defined the boundary conditions of *f* = 1 for the inner surface and *f* = 2 for the pial surface, we then solved Laplace’s equation to get a smooth transition of values from the white matter surface to the pial surface within the cortical mask of the whole 3D volume, where the gradient ∇(*x, y, z*) at each voxel can be calculated.

For the streamlined calculation, we used a previously estimated mid-surface for generating starting points. We took the mid-surface from our BMA2019 flat map package and applied the transformation from BMA2019 to BMA2.0 to transform all vertices on the mid-surface to BMA2.0 space.

All 20 cortical parcellations were loaded (10 original and 10 flipped), and streamline starting points were generated by first identifying 3D vertices of the mid-cortical and verifying their overlap with the BMA2.0 cortical mask. Four random points per mid-surface triangle were generated, checked to be within the cortical mask, and added to the list, increasing the number of starting points to 1,336,640. The gradient at each voxel of the Laplace volume was calculated to provide the direction for each streamline to follow in 3D. From the starting point, a streamline followed the gradient outwards in two directions towards the white matter and pial surfaces of the cerebral cortex. At each step, label assignments were collected from all 20 cortical parcellations along the streamline until it reached the white matter or pial surface. Finally, the label with the highest probability along the streamline was selected and assigned to all voxels it intersected with. This was carried out for all streamlines to create a final population-based parcellation in the standard space.

#### Subcortical structure delineation

Subcortical regions for our BMA2.0 region parcellations were based on a previous release (BMA2017), where the subcortical regions were delineated by Hashikawa et al.^18^ based on a single brain. This was a compromise due to the amount of additional work required to annotate the subcortical regions for 10 individuals within a reasonable time frame. For image registration, the BMA2017 Nissl brain was initially registered to the BMA2.0 average Nissl template, incorporating MRI contrast as a constraint, as the regions were identified using Nissl contrast in BMA2017. Next, we used the transforms from registration results to transform the BMA2017 labels into the BMA2.0 space. We then used the cortical mask to exclude the cortical regions and added the remaining subcortical regions into our BMA2.0 region parcellations.

Using 3D Slicer, the subcortical region labels were manually edited to improve the alignment to our population average template. We prioritized misalignments that were visible and easily correctable. Subcortical regions with misalignments were isolated based on their region label number. These were then manually edited slice by slice across the entire brain, using all three contrasts (MRI, myelin, and Nissl) with the original BMA2017 serving as a reference to guide our manual segmentation, as some subcortical region borders can only be identified in one contrast. After each region edit, we inspect them in all three views (axial, coronal, and sagittal) and apply 3D smoothing. A total of 50 subcortical regions were refined in this manner.

In comparison to the original BMA2017 subcortical parcellations, we combined several subregions within the inferior olive (IC), namely, Inferior olive; dorsal nucleus (IOD) and Inferior olive; principal nucleus (IOPr) as we could not correctly delineate their boundaries using our average template contrasts.

#### Cerebellar lobule delineation

To achieve accurate anatomical delineation of the cerebellum for the BMA2.0, we manually segmented each lobule, resulting in the identification of 45 distinct cerebellar cortical regions and refined four deep cerebellar nuclei (DCN). Each cerebellar lobule was annotated in the sagittal plane, which provided optimal visualization of lobular boundaries. At the same time, the DCN were delineated in the coronal plane to maximize clarity of internal nuclear structures.

These structures were manually annotated using high-resolution (40 µm) T2WI ex vivo MRI in conjunction with corresponding Nissl-stained histological sections. The delineation process was cross-validated with the reference atlas provided by Lin et al.^44^, both through online access and in-person consultation.

To ensure accurate boundary demarcation between the white matter and the granular layer (GL), we additionally performed manual segmentation of the arbor vitae. This characteristic, tree-like branching structure of the cerebellar white matter, was essential for precisely refining lobular borders within the BMA2.0 ex vivo T2WI population average template.

We refined the cerebellar lobule definitions by comparison with the Lin et al.^44^ reference, ultimately establishing a total of 45 cerebellar regions per hemisphere. The incorporation of these detailed cerebellar region parcellations significantly enhances the anatomical completeness of BMA2.0, supporting comprehensive studies of whole-brain functional networks, structural connectivity, and regional integration across the marmoset brain.

### Flat map and cortical surfaces

For the flat map and cortical surfaces, we used ANTs transforms from BMA2019 to BMA2.0 to transform the vertex coordinates of the cortical surfaces from the BMA2019 space to the BMA2.0 space, thereby obtaining cortical surfaces in the BMA2.0 template shape. The original cortical surfaces were created based on a mid-cortical thickness boundary in the BMA2019 T2WI template space, using the Caret software by Van Essen et al.^45^. The white matter and pial surfaces were generated from this mid-thickness surface by starting at the vertices of the surface and following the gradient of the Laplace’s equation solution (for BMA2019’s cortical region with boundary conditions defined at the white matter and pial boundaries) until the cortical boundaries were reached. We used the command line program (wb_command -volume-to-surface-mapping) from Connectome Workbench^46^ to map the BMA2.0 region labels to the transformed surfaces, providing us with region labels for coordinate mapping on the flat map. The surfaces and flat maps for both hemispheres are shown in Figure 1D and E.

### Interoperability

As part of the BMA2.0 package, we have included both forward and backward transforms in our previous releases, BMA2017 and BMA2019. For BMA2017, we used cortical and background segmentation as constraints to register the BMA2017 T2WI MRI with the BMA2.0 T2WI average template. Similarly, segmentation constraints were used to accurately register between the BMA2019 T2WI average template and the BMA2.0 T2WI average template. These transforms can be used to map any data that was registered to BMA2017 or BMA2019 to our new BMA2.0 space, including mapping to the BMA2.0 cortical surfaces and flat maps. To further expand the interoperability of BMA2.0, we also provide transformations to other widely used marmoset brain atlases, namely the Marmoset Brain Mapping Atlas Version 2 (MBMv2)^5^, the Marmoset Brain Connectivity Atlas (MBC)^47^, and the Subcortical Atlas of the Marmoset (SAM)^28^. These transformations enable users to map data between BMA2.0 and these spaces, taking advantage of additional data, such as the white matter pathways from MBMv2 or the tracer projections from the Marmoset Brain Connectivity Atlas. We have also included the white matter parcellations of Liu et al.^5^, mapped to the BMA2.0 standard space, in our dataset. This provides complete coverage of both gray and white matter structures when using our atlas.

This interoperability enables comparative studies and the integration of datasets generated using different reference frameworks, thereby enhancing the accessibility of BMA2.0 for previous studies.

## Data Records

The BMA2.0 dataset consists of three main components. Firstly, the atlas component of the dataset includes volumetric files in compressed NIfTI format (.nii.gz) in 100 µm resolution, includes region labels and multimodal templates. The corresponding region names and color lookup table are provided in text format (.ctbl). A resampled 50 µm resolution version of the region labels is also included for better visualization. Next, the flat map component of the dataset consists of the flat map, region labels, and cortical surfaces (pial, mid-thickness, and white matter) for both hemispheres, in GIfTI format (.gii). Descriptions of the region parcellations and templates are given in Table 2, and the flat map and cortical surfaces files are described in Table 4. Lastly, the transformations component of the dataset includes forward and inverse warp fields in compressed NIfTI format (.nii.gz) and affine transforms (.mat), providing the non linear and linear transform components between BMA2.0 and other template spaces (BMA2017^16^, BMA2019^17^, MBMv2^5^, MBC^47^, and SAM^28^). The white matter parcellations of MBMv2^5^ mapped to BMA2.0 are detailed in Table 3. The transformation files are detailed in Table 5.

**Table 2.** Data records of BMA2.0 atlas and templates.

| Filename | File type | Description |
| --- | --- | --- |
| BMA2.0_avg_exvivo_T2WI.nii.gz | NIFTI | Population average ex vivo T2WI MRI template ($N = 91$) |
| BMA2.0_avg_invivo_T2WI.nii.gz | NIFTI | Population average in vivo T2WI MRI template ($N = 446$) |
| BMA2.0_avg_myelin.nii.gz | NIFTI | Population average myelin contrast template ($N = 10$) |
| BMA2.0_avg_nissl.nii.gz | NIFTI | Population average Nissl contrast template ($N = 10$) |
| BMA2.0_regions_label.nii.gz | NIFTI | BMA2.0 region labels |
| BMA2.0_regions_label_50mu.nii.gz | NIFTI | BMA2.0 region labels (resampled to 50 $\mu\text{m}$) |
| BMA2.0_regions_list.ctbl | Text | Region and color lookup table for BMA2.0 atlas |
| BMA2.0_slicer_scene.mrml | XML | 3D Slicer scene file for BMA2.0 atlas and templates |

**Table 3.** Data records of MBMv2 white matter parcellations mapped to BMA2.0.

| Filename | File type | Description |
| --- | --- | --- |
| MBMv2_wm_ROIs_1_to_BMA2.0.nii.gz | NIFTI | MBMv2 white matter ROI 1 mapped to BMA 2.0 |
| MBMv2_wm_ROIs_2_to_BMA2.0.nii.gz | NIFTI | MBMv2 white matter ROI 2 mapped to BMA 2.0 |
| MBMv2_wm_ROIs_3_to_BMA2.0.nii.gz | NIFTI | MBMv2 white matter ROI 3 mapped to BMA 2.0 |
| MBMv2_wm_ROIs_1_labels.ctbl | Text | Region and color lookup table for MBMv2 white matter ROI 1 |
| MBMv2_wm_ROIs_2_labels.ctbl | Text | Region and color lookup table for MBMv2 white matter ROI 2 |
| MBMv2_wm_ROIs_3_labels.ctbl | Text | Region and color lookup table for MBMv2 white matter ROI 3 |

**Table 4.**
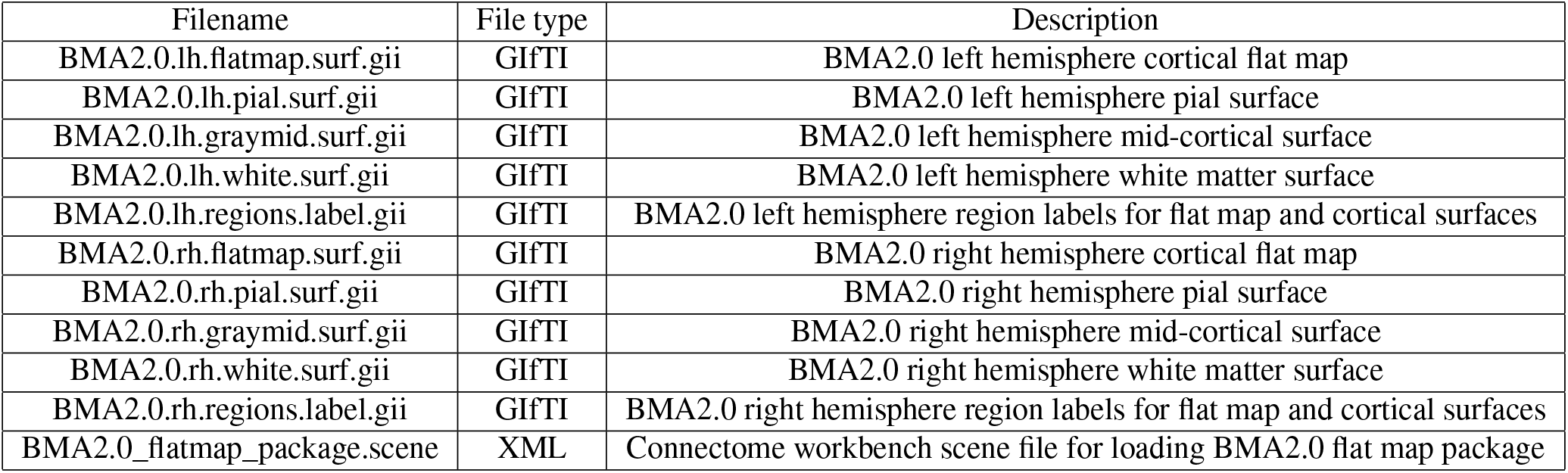
Data records of BMA2.0 cortical flat maps and surfaces.

**Table 5.**
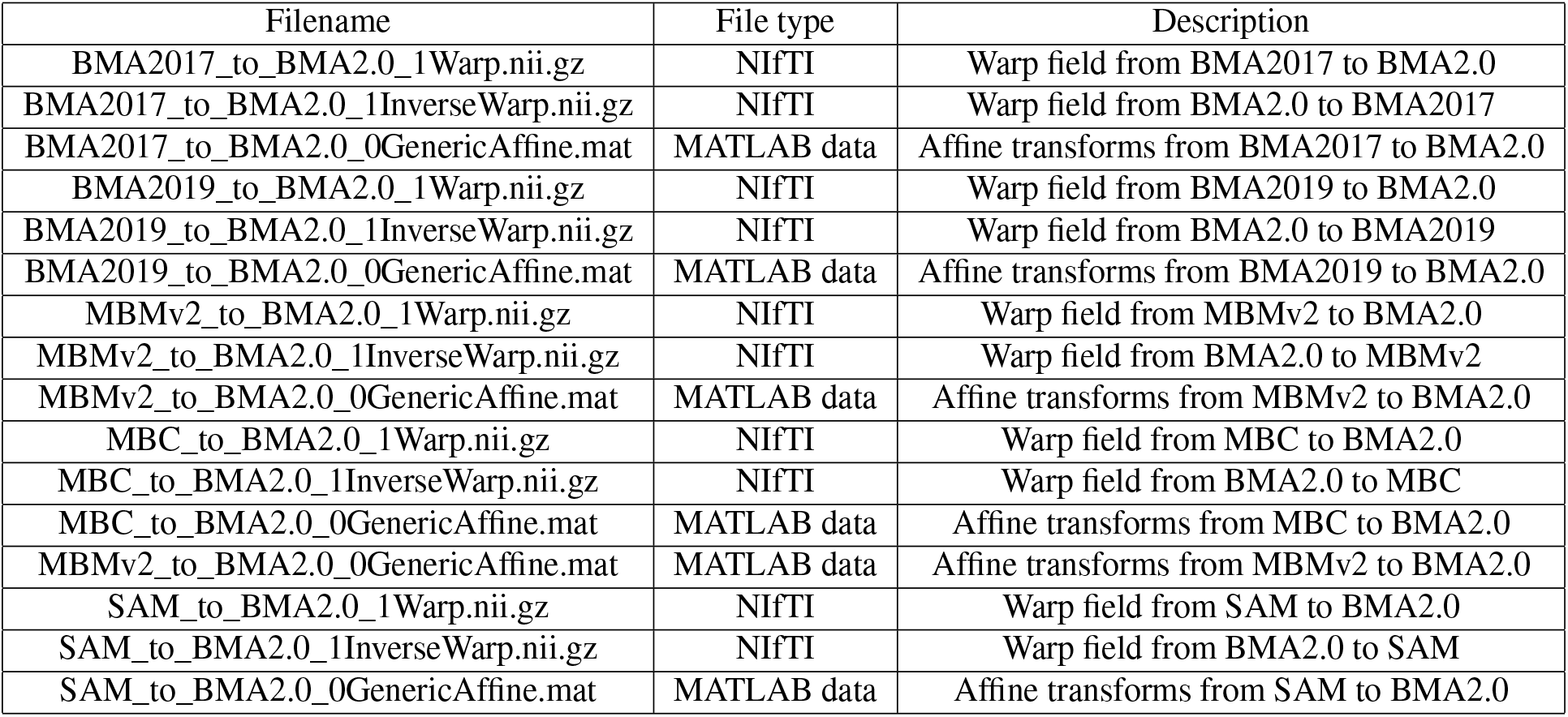
Data records of transformations (affine and warp fields) between BMA2.0 and other marmoset brain atlas templates.

| Filename | File type | Description |
| --- | --- | --- |
| BMA2017_to_BMA2.0_1Warp.nii.gz | NIfTI | Warp field from BMA2017 to BMA2.0 |
| BMA2017_to_BMA2.0_1InverseWarp.nii.gz | NIfTI | Warp field from BMA2.0 to BMA2017 |
| BMA2017_to_BMA2.0_0GenericAffine.mat | MATLAB data | Affine transforms from BMA2017 to BMA2.0 |
| BMA2019_to_BMA2.0_1Warp.nii.gz | NIfTI | Warp field from BMA2019 to BMA2.0 |
| BMA2019_to_BMA2.0_1InverseWarp.nii.gz | NIfTI | Warp field from BMA2.0 to BMA2019 |
| BMA2019_to_BMA2.0_0GenericAffine.mat | MATLAB data | Affine transforms from BMA2019 to BMA2.0 |
| MBMv2_to_BMA2.0_1Warp.nii.gz | NIfTI | Warp field from MBMv2 to BMA2.0 |
| MBMv2_to_BMA2.0_1InverseWarp.nii.gz | NIfTI | Warp field from BMA2.0 to MBMv2 |
| MBMv2_to_BMA2.0_0GenericAffine.mat | MATLAB data | Affine transforms from MBMv2 to BMA2.0 |
| MBC_to_BMA2.0_1Warp.nii.gz | NIfTI | Warp field from MBC to BMA2.0 |
| MBC_to_BMA2.0_1InverseWarp.nii.gz | NIfTI | Warp field from BMA2.0 to MBC |
| MBC_to_BMA2.0_0GenericAffine.mat | MATLAB data | Affine transforms from MBC to BMA2.0 |
| MBMv2_to_BMA2.0_0GenericAffine.mat | MATLAB data | Affine transforms from MBMv2 to BMA2.0 |
| SAM_to_BMA2.0_1Warp.nii.gz | NIfTI | Warp field from SAM to BMA2.0 |
| SAM_to_BMA2.0_1InverseWarp.nii.gz | NIfTI | Warp field from BMA2.0 to SAM |
| SAM_to_BMA2.0_0GenericAffine.mat | MATLAB data | Affine transforms from SAM to BMA2.0 |

BMA2.0 comprises a total of 323 region parcellations per hemisphere, consisting of population-based cerebral cortex delineations (*N* = 10, with an effective sample size of 20 cortex delineations due to symmetry) of 117 regions, 156 subcortical parcellations, and 45 cerebellar lobule parcellations.

The region name and color lookup table for the cerebral cortex and subcortical areas were adopted from BMA2017^16^, which was first defined by Hashikawa et al.^18^. We have extended the list by adding 45 cerebellar lobule delineations per hemisphere (region ID 787-831 on the right hemisphere and 10787-10831 on the left hemisphere). Cerebellar region name and color were defined by Lin et al.^44^, as shown in Figure 7A.

**Figure 6.**
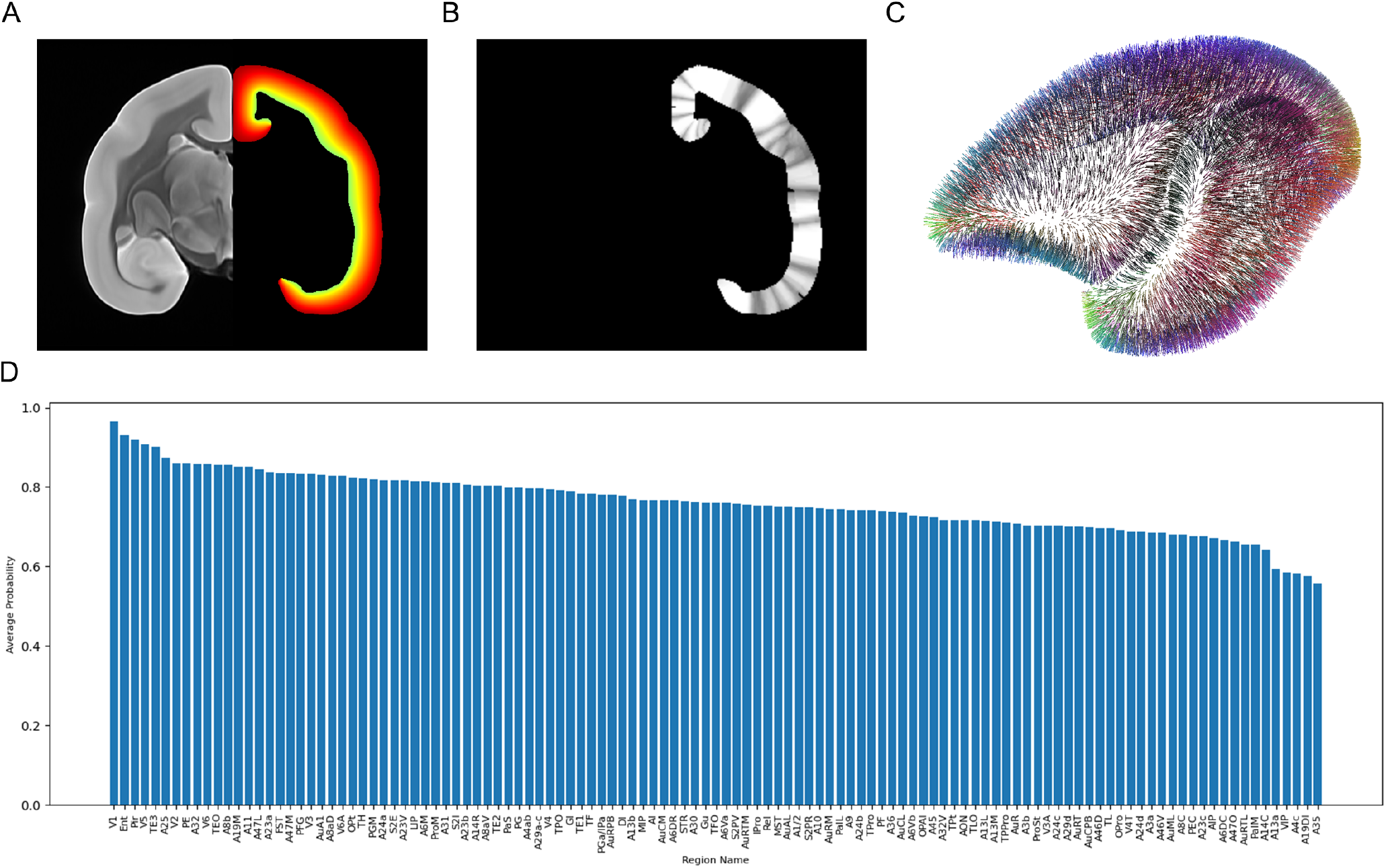
Streamline-based label assignment using the Laplace approach. (A) A coronal slice of the Laplace’s equation solution volume for the left hemisphere using boundary conditions defined at the white matter surface (maximally yellow) and the pial surface (maximally orange). The right hemisphere shows the average MRI template for reference. (B) Visualization of label assignment probabilities within the cerebral cortex after streamline calculation for a coronal slice; lighter gray indicates higher probability and darker gray lower. (C) 3D visualization of streamlines generated from the cortical mid-thickness surface towards the pial surface, verifying streamline correctness. (D) Graph of average label assignment probabilities per label, sorted from highest to lowest probability.

**Figure 7.**
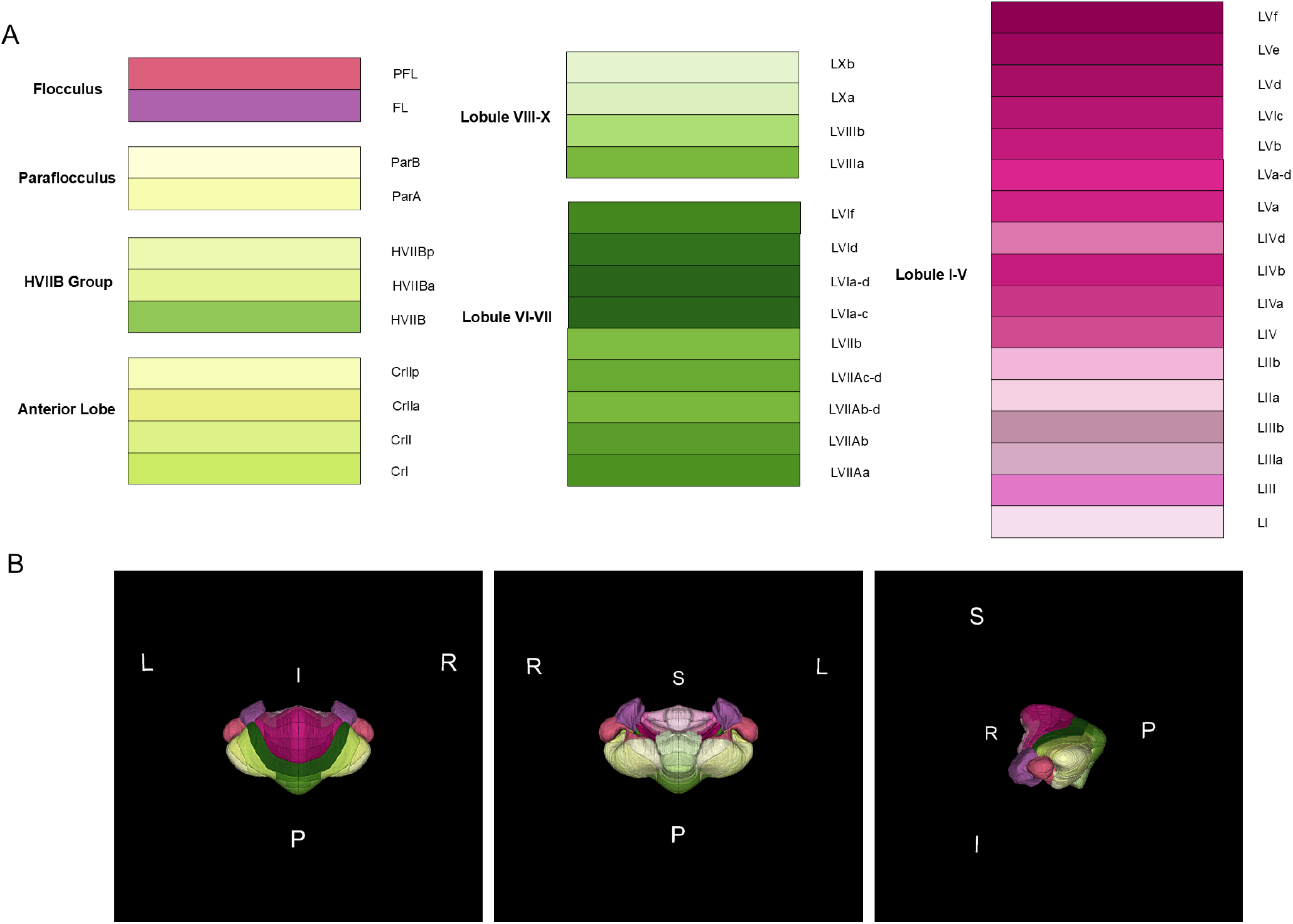
Overview of the cerebellum region parcellations. (A) Color coding table of the cerebellum region parcellations by Lin et al.^44^, with region groups representing main brain function. (B) 3D reconstruction of the cerebellum region parcellations in superior, inferior, and left hemisphere views.

## Technical Validation

### Confidence of streamline-based label assignment

During streamline calculation, we collected data on the confidence of each label assignment. Figure 6D shows an example visualization of this data in the coronal view, with darker shades of gray indicating lower confidence and lighter shades of gray indicating higher confidence. A more detailed analysis of the average label assignment confidence is shown in Figure 6E, which shows the average confidence of label assignments for each cortical region. Here, we were expecting the confidence to be low for regions around the lateral sulcus, due to individual variability and the difficulty of image registration in that area, but we were surprised to find out that the bottom 5 regions with the lowest confidence were Area 35 of cortex (A35), Area 19 of cortex; dorsointermediate part (A19DI), Area 4 of cortex; part c (A4c), Ventral intraparietal area of cortex (VIP), and Area 13a of cortex (A13a). These are smaller regions that are not around the lateral sulcus. This result can be attributed to a greater overlap occurring between these regions and their surrounding larger regions. From Figure 6E, we can see that even the lowest average probability is close to 0.6, showing that the majority of assignments to a region are consistently occurring.

### Regional variation in supragranular layer thickness and system-level gradients

To quantify the thickness of supragranular layers (layers 1-3) across the marmoset neocortex, we used the population average myelin-stained volume from BMA2.0. Myelin staining exhibits a robust laminar contrast in the form of a prominent “middle band,” which corresponds approximately to the layer 3-4 boundary. Compared to Nissl staining, which requires high-magnification inspection and is less reliable at lower resolution due to variability in cell density and staining intensity, this feature provides a more consistent and reproducible laminar landmark across regions and individuals.

The upper boundary of the middle band was manually traced by a well-trained neuroanatomist (N.I.) based on histological expertise. For each cortical location, the distance from the pial surface to the traced upper edge of the middle band was computed using a Laplace equation streamline coordinate system defined between the pial and white matter surfaces. The resulting measurement was used as a proxy for the combined thickness of layers 1-3. These thickness values were projected onto the 2D flat map representation of the BMA2.0 atlas to visualize spatial distributions. The resulting maps revealed systematic gradients along dorsal-ventral and anterior-posterior axes, consistent with previously described anatomical and functional hierarchies in the primate cortex (Figure 8B, C). Association areas, such as TE and ProM, exhibited the largest supragranular thickness, whereas motor areas and medial wall regions (e.g., A4ab, A6M) consistently showed thinner upper layers.

**Figure 8.**
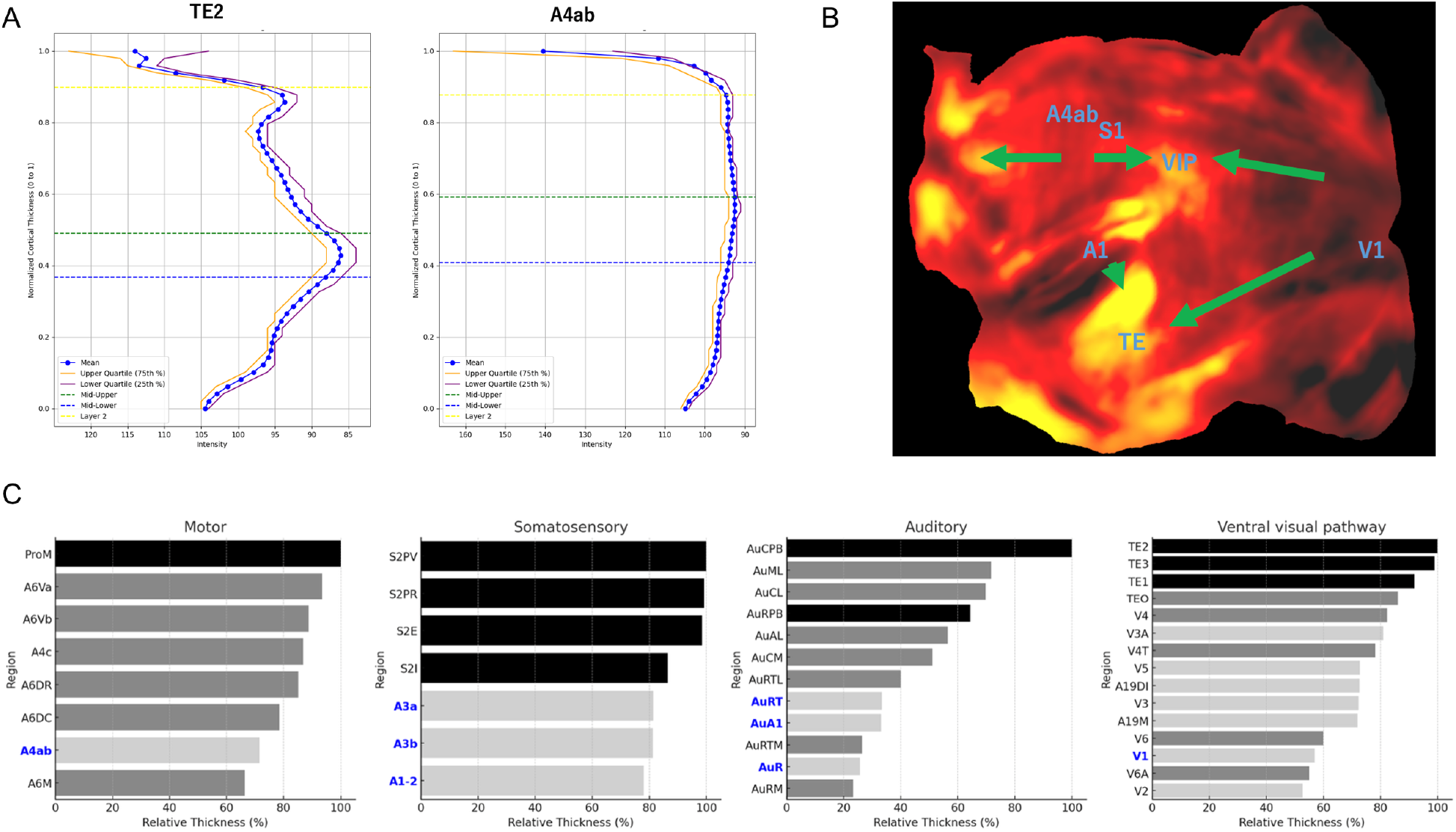
Regional gradients and cytoarchitectonic features of supragranular (layer 1-3) thickness across the marmoset cortex. (A) Nissl intensity profiles along normalized cortical depth (from pial = 0 to white matter = 1) for two representative areas: TE2 (temporal association cortex) and A4ab (primary motor cortex). TE2 shows a prominent peaked middle band (of lower intensity values) associated with a dense Nissl profile at mid-depth. In contrast, A4ab—an granular area—shows a flat Nissl profile with no clear middle band, consistent with its lack of layer 4. (B) Thickness of supragranular layers (L1-3) across the neocortex, computed as the distance from the pial surface to the upper edge of the myelin-defined middle band. Values are projected onto the BMA2.0 flat map. Warmer colors indicate greater thickness. Arrows highlight example regions from different cortical systems. Arrows indicate presumed hierarchical progression within each cortical system (motor, somatosensory, auditory, and visual), anticipating the quantitative gradients shown in Panel C. (C) Bar plots showing normalized supragranular (L1-3) thickness across four cortical systems. Motor system: primary motor cortex (A4ab) →premotor areas (A6) →promotor (ProM). Somatosensory system: primary somatosensory areas (A3a, A3b, A1/2) →secondary somatosensory areas (S2). Auditory system: core regions (AuA1, AuR, AuRT) →belt and parabelt regions. Ventral visual stream: V1 →V2-V4 →inferotemporal areas (TE). Within each system, L1-3 thickness increases along presumed cortical hierarchies, supporting the idea that supragranular expansion correlates with integrative processing demands. Gray scale indicates hierarchical level (black: higher, dark gray: middle, light gray: lower), and primary areas are highlighted in blue.

To interpret these patterns in relation to underlying cytoarchitecture, we extracted Nissl intensity profiles across normalized cortical depth for representative areas. In TE2, a distinct contrast corresponding to the middle band was observed, whereas in A4ab, the middle band was weak or absent (Figure 8A). This finding is consistent with the classical notion that A4ab is an agranular region, lacking a well-defined layer 4. Moreover, in TE2, the Nissl density peak corresponded closely with the location of the myelin-defined middle band, suggesting that in this region, the middle band reflects the anatomical position of layer 4.

We further compared supragranular thickness across major functional systems using representative regions (Figure 8C). In the motor-frontal system, a rostro-caudal gradient was observed, with ProM and A6Va exhibiting thicker upper layers than A4ab. In the somatosensory system, secondary and belt areas such as S2PV and S2PR had greater L1-3 thickness than primary sensory areas like A1/2. The auditory system exhibited increasing supragranular thickness from the core to the belt and parabelt regions. Similarly, in the ventral visual stream, TE2 exhibited the largest thickness of L1-3, with a progressive reduction toward V1.

Comparative analysis across cortical systems (Figure 8B, C) demonstrated that supragranular thickness increases systematically along putative processing hierarchies—for example, from V1 to TE2 in the ventral visual stream, and from primary to higher regions in the motor, somatosensory, and auditory cortices (Figure 8B, C). These findings support the idea that supragranular expansion is a structural correlate of integrative processing and cortical complexity.

Taken together, these results demonstrate that the BMA2.0 atlas provides not only an anatomical coordinate framework but also a morphometric substrate for studying cortical microstructure. This approach may further facilitate future efforts to build probabilistic laminar maps across individuals and species, analogous to developments in rodent and human brain mapping.

### Structural and functional comparison by MTESS

We investigated whether the BMA2.0 anatomical segmentation reflects functional activity. To do this, we used awake resting-state (rs-) fMRI data from four marmosets^48,49^ and quantified the similarity of multivariate time-series across 12 sessions within each individual using the Multivariate Time-series Ensemble Similarity Score (MTESS)^50^. We hypothesized that if the region segmentation reflects functional activity, the similarity of multivariate time-series within an individual would be higher than that using non-functional segmentation. Transforming the BMA2.0 atlas into the rs-fMRI image space resulted in 544 ROIs (Figure 9A top). In addition, K-means clustering was performed based on voxel distance to divide the marmoset brain into 544 regions, named DistKm (Figure 9A bottom). DistKm does not follow an anatomical regional division at all. By comparing the similarity of within-individual multivariate time-series using the ROIs of BMA2.0 and DistKm, it is possible to test whether anatomical regional division has no functional meaning (the null hypothesis). We calculated the MTESS for each session (resulting in a 48 × 48 matrix) for the four marmosets (Figure 9B), extracted within-individual session comparisons, and compared BMA2.0 with DistKm (Figure 9C). Results showed that BMA2.0 had significantly higher similarity, better capturing functional activity, and suggesting that anatomical segmentation has functional meaning.

**Figure 9.**
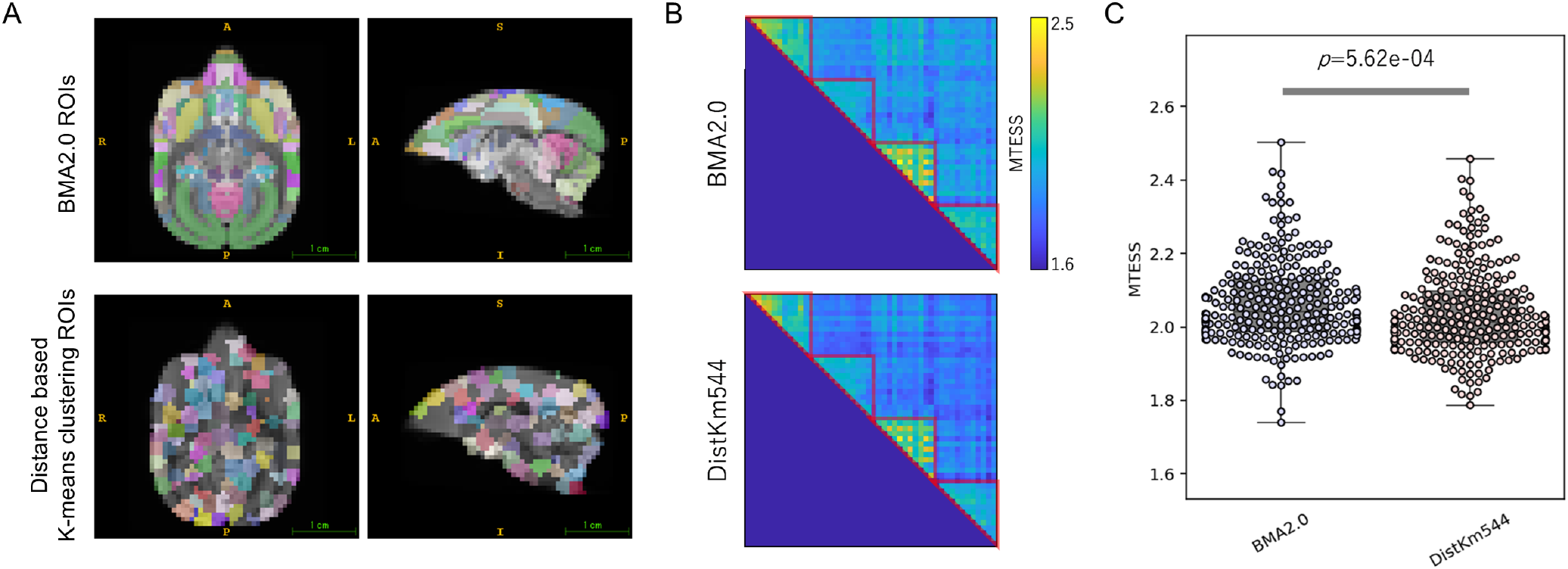
Comparing multivariate time-series similarity between BMA2.0 and distance-based K-means clustering ROIs. (A) Example horizontal (*z* = 22*/*45) and sagittal (*x* = 22*/*45) planes in each ROI type. (top) BMA2.0 atlas ROIs. (bottom) distance-based K-means (DistKm) clustering ROIs. (B) MTESS matrix results of BMA2.0 ROIs (top) and DistKm ROIs (bottom). Similarity of rs-fMRI multivariate time-series of 4 animals x 12 sessions (48 sessions in total). The higher the value, the greater the similarity between the multivariate time series across sessions. Red triangles indicate inter-session similarity for each individual. (C) Similarity between individual sessions for each ROI type (red triangles in B); BMA2.0 atlas is significantly higher than DistKm ROIs (*p* = 5.62*e*− 04, Mann-Whitney U test).

### Connectomics using the BMA2.0 in vivo average template

To demonstrate the applicability of our BMA2.0 atlas, we used the in vivo population template and created a population average in vivo Diffusion weighted Imaging (DWI) volume based on the NA216 dataset (*N* = 126, 96-channel NIfTI volumes) by Hata et al.^21^. These were used to perform fiber tracking and generate a population-based connectome of the common marmoset brain. The b0 contrast of each scan was then registered to the BMA2.0 in vivo average template using ANTs. The resulting transform files for each brain were then used to transform all separated 96 channels of the individual dataset to the BMA2.0 average space. Averages for each of the 96 channels, with 126 datasets per channel, were calculated. The 96 averaged channels were then reconstructed into a single NIfTI file, yielding a population-average DWI.

Fiber tracking was performed using DSI Studio’s^5^ tractography tool^51–54^ using generalized q-sampling imaging (GQI) on the calculated population average DWI and its b-value and b-vectors. 5,000,000 tracks were deterministically generated. The fiber tracking result is shown in Figure 10A, overlaid with the BMA2.0 in vivo average template, and the fiber density image shown in B. The average tract length was 24.75 mm, consistent with another marmoset DWI tractography study by Schaeffer et al.^55^.

**Figure 10.**
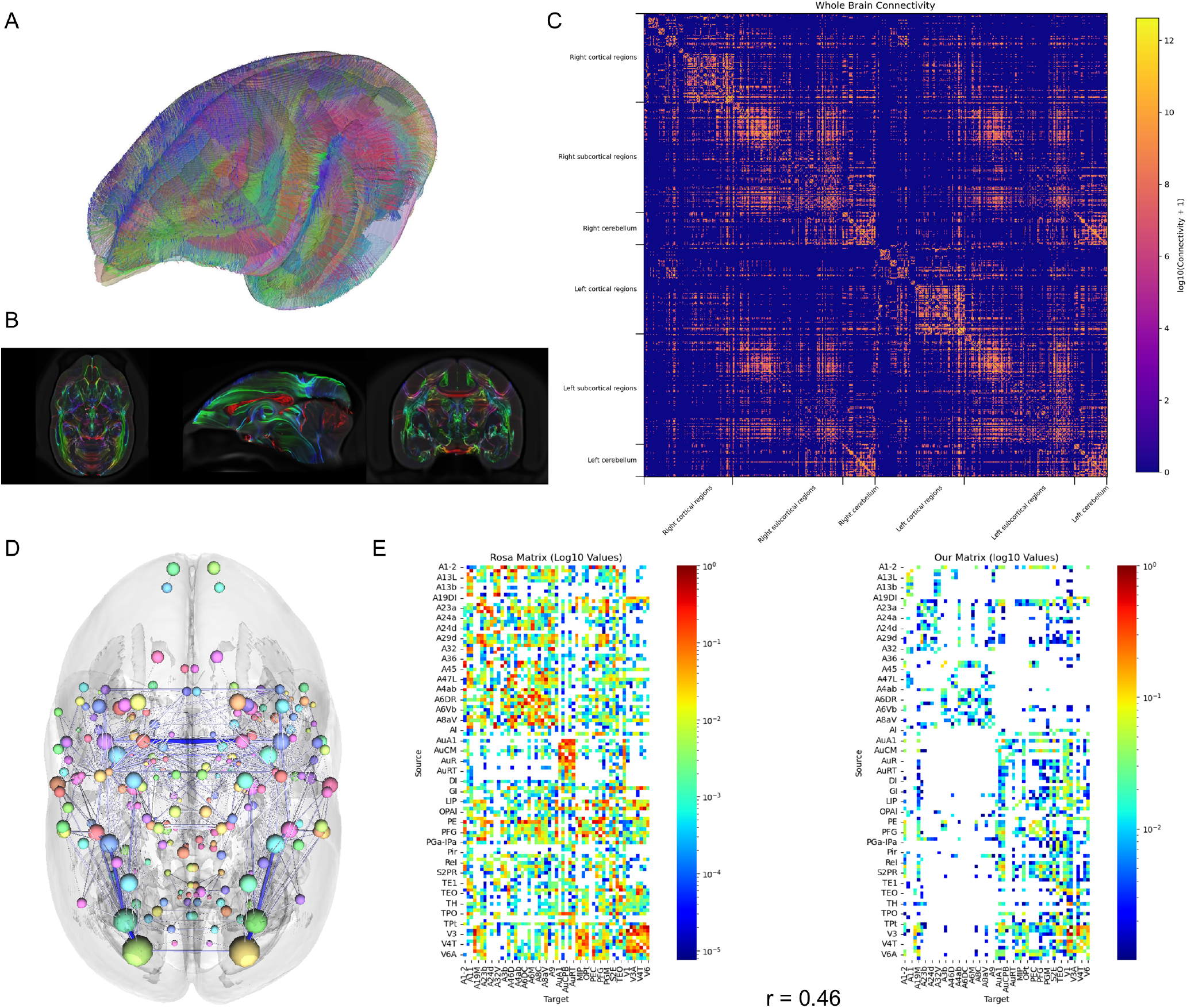
Population-based tractography calculated using the BMA2.0 in vivo average template, and analysis of the connectivity matrix generated from tractography. (A) Visualization of fiber tracking (DSI Studio) using the population average in vivo diffusion MRI data (*N* = 126) with BMA2.0 whole brain regions overlay (*ROI* = 636). (B) Axial, sagittal, and coronal views of fiber tracking density visualized in diffusion space overlayed onto BMA2.0 in vivo average template. (C) Whole brain connectivity matrix (*ROI* = 636) generated from fiber tracking using the in vivo population average DWI data. (D) 3D visualization of region connectivity in axial view, showing strong connections between posterior visual regions. (E) Comparison of connectivity matrices between the marmoset brain connectivity atlas project^47^ and the matrix calculated in the present work, using the same ROIs and region ordering. The Pearson correlation score between the two matrices was 0.46.

The population average connectome of the marmoset brain was calculated using the BMA2.0 regions (*N* = 636) with fiber tracking information. The result is shown in Figure 10C, which shows the strength of connection between all brain regions.

To assess the anatomical validity of the BMA2.0 diffusion-based connectivity matrix, we compared it with the axonal tracer-based interareal connectivity matrix reported by Majka et al.^47^, which is one of the most comprehensive mesoscale cortical connectivity maps available for the marmoset. We selected corresponding cortical regions and ordered the data to simplify the comparison – see Figure 10E. Despite being derived from entirely different individuals and using various methodologies (in vivo DWI vs. retrograde tracer injections), we observed a substantial correlation (Pearson’s *r* = 0.4605) between the diffusion tractography-based and tracer-based area-to-area connectivity weights. This degree of correspondence is comparable to previous large-scale cross-modal studies in mice, such as the Allen Mouse Brain Connectivity Atlas (Spearman r ≈0.46 for area level comparisons), and underscores the utility of the BMA2.0 connectome for network-level analyses. For non-human primates, a landmark study using the same individual macaque for both tracer injections and postmortem DWI reported a higher correlation (*r* ≈ 0.56) under idealized conditions (Donahue et al.^56^). However, such designs are logistically constrained and not generalizable. Our cross-individual result (*r* = 0.4605) thus demonstrates a high degree of robustness and validates the anatomical relevance of our connectome under realistic application conditions.

Given the challenging nature of intermodal normalization, especially in primates where individual anatomical variation is high, this level of agreement highlights the potential of BMA2.0 to serve as a common coordinate framework for integrative studies. We expect that such validated connectomic maps will enhance downstream analyses, including computational modeling, cross-species comparisons, and integration with functional and genetic data.

## Usage Notes

All NIfTI data provided in the BMA2.0 atlas package are compatible with a wide range of neuroinformatics and image analysis software. These files can be loaded using software such as 3D Slicer^38^ (version 5.6.2) and ITK-SNAP^57^ (version 4.0.2), allowing users to inspect and explore BMA2.0 templates alongside the atlas. We have provided a 3D Slicer scene file (.mrml) that the user can open, which loads all the templates, region labels, and region definitions of BMA2.0.

We recommend using Python for code-based processing of NIfTI files for BMA2.0. Python libraries such as Nibabel^58^ (version 5.3.2) and SimpleITK^59^ (version 2.5.0) provide access to both voxel data and metadata, which contain information on resolution, voxel spacing, orientation, etc.

Registering new data to our multimodal templates can be done using ANTs’ command line interface. A tutorial on how to perform image registration and apply transforms using ANTs can be found in the Brain/MINDS data portal^6^. For interpolations, users should use “NearestNeighbor” when transforming labels and “Linear” for templates.

For users who prefer working with Python, we recommend using the ANTsPy^31^ library, which provides a Python interface to ANTs, offering equivalent functionality such as reading NIfTI files, performing image registration, and applying transformations.

The flat map and cortical surfaces can be loaded and visualized using Connectome Workbench^46^ (version 2.1.0). We provide a scene file (.scene) for Connectome Workbench that will load the flat map, cortical surfaces, and labels. For users who would like to access flat map and surface data using Python, we recommend using the Nibabel^58^ (version 5.3.2) library.

## Data Availability

The BMA2.0 atlas package includes the region parcellations, multimodal templates, cortical flat map and surfaces, and forward and backward transforms to other atlases. It is publicly available on Figshare: [The Brain/MINDS 3D Digital Marmoset Brain Atlas Version 2.0](https://doi.org/10.6084/m9.figshare.29992687).

## Code Availability

Image data processing and region parcellation construction relied on a combination of custom Python scripts and extensive manual processing. These scripts are not sufficiently documented for general use and are therefore not publicly released. All data processing can be reproduced by following the procedures and software packages described in the Methods section. Additional details about the region parcellations and templates construction can be provided by contacting the corresponding author.

## Acknowledgements

This research was supported by Japan Agency for Medical Research and Development (AMED) under Grant Numbers JP15dm0207001 and JP23wm0625001. This work was supported by JSPS KAKENHI under Grant Numbers JP22H05163 and 24K03256 to K.N. This research was supported by Japan Agency for Medical Research and Development (AMED) under Grant Numbers: JP19dm0207088, JP24wm0625414, and JP24wm0625404 to K.N

## Author Contributions Statement

A.W formulated the project. R.G and A.W developed the workflow and processing pipelines, performed all the data processing to construct the region parcellations, multimodal templates, transform data, flat map, and their corresponding surfaces. N.I, H.A and T.T collected the raw data, performed histological staining, and delineated cortical regions. K.N provided region segmentation data for T2WI MRI data. J.H provided T2WI MRI data for in vivo and ex vivo template construction. R.G, K.N, S.H and J.W edited subcortical regions supervised by N.I. M.L performed cerebellar lobule delineation. R.G and N.I analyzed DWI-based tractography and estimation of layer 1-3 thickness using myelin contrast. T.O performed functional dynamics comparison by MTESS. S.I, P.D, T.Y, and H.O contributed to the review and editing of the manuscript. All authors reviewed the manuscript.

## Footnotes

1 https://dataportal.brainminds.jp

2 https://github.com/ANTsX/ANTs

3 See https://dataportal.brainminds.jp/tutorial-how-to-make-a-symmetric-brain for a detailed methodology

4 https://www.slicer.org/

5 https://dsi-studio.labsolver.org/

6 https://dataportal.brainminds.jp/ants-tutorial

